# Human antibodies against West Nile and related orthoflaviviruses

**DOI:** 10.64898/2026.04.02.715800

**Authors:** Tomás Cervantes Rincón, Tereza Frckova, Zaria I. Contejean, Jasmine Cantergiani, Kevin Groen, Benedetta Cena, Simone G. Moro, Filippo Bianchini, Luca Simonelli, David Jarrossay, Silvia Tosolini, Roger Kuratli, Anna R. E. Robinson, Monika Cizkova, Emily G. Niejadlik, Jacques Moritz, Roshan Thakur, Zuzana Krátka, Dragana Mijatović, Jasmina Grujić, Jiri Holoubek, Zorana Budakov-Obradović, Jiri Salat, Václav Hönig, Marija Vraneš, Zvezdana Lojpur, Dajana Lendak, Siniša Sević, Monika Bajči, Lidija Popović-Dragonjić, Biljana Popovska Jovicic, Jagoda Gavrilovic, Tania Kapoor, Margaret R. MacDonald, Stylianos Bournazos, Luca Varani, Martin Palus, Benjamin G. Hale, Pavle Banović, Daniel Ruzek, Christopher O. Barnes, Davide F. Robbiani

**Affiliations:** Institute for Research in Biomedicine, Università della Svizzera italiana, 6500 Bellinzona, Switzerland; Institute of Experimental Biology, Faculty of Science, Masaryk University, Brno, Czech Republic; Veterinary Research Institute, Brno, Czech Republic; Stanford Biosciences, Stanford School of Medicine, Stanford, CA 94305, USA; Institute of Medical Virology, University of Zurich, 8057 Zurich, Switzerland; Institute of Parasitology, Biology Centre of the Czech Academy of Sciences, Ceske Budejovice, Czech Republic; Faculty of Science, University of South Bohemia, Ceske Budejovice, Czech Republic; The Rockefeller University, Laboratory of Molecular Genetics and Immunology, New York, NY 10065, USA; Diagnostics and Laboratory Research Task Force, Balkan Association for Vector-Borne Diseases, 21000 Novi Sad, Serbia; Department for Research & Monitoring of Rabies & Other Zoonoses, Pasteur Institute Novi Sad, 21000 Novi Sad, Serbia; Blood Transfusion Institute Vojvodina, 21000 Novi Sad, Serbia; Faculty of Medicine in Novi Sad, University of Novi Sad, 21000 Novi Sad, Serbia; Department for Transfusion Medicine, Clinical Center of Kragujevac, Kragujevac, Serbia; University of Kragujevac, Faculty of Medical Sciences, Serbia; Clinic for Infectious Diseases, Clinical Center of Vojvodina, 21000 Novi Sad, Serbia; Department of Infectious Diseases and Epidemiology, Faculty of Medicine Niš, University of Niš, 18000 Niš, Serbia; Clinic for Infectology, University Clinical Center Niš, 18000 Niš, Serbia; Clinical medicine Task Force, Balkan Association for Vector-Borne Diseases, 21000 Novi Sad, Serbia; University of Kragujevac, Faculty of Medical Sciences, Department of Infectious Diseases, Serbia; University Clinical Centre Kragujevac, Serbia; The Rockefeller University, Laboratory of Molecular Immunology, New York, NY 10065, USA; The Rockefeller University, Laboratory of Virology and Infectious Disease, New York, NY 10065, USA; Department of Prevention of Rabies and Other Infectious Diseases, Pasteur Institute Novi Sad, 21000 Novi Sad, Serbia; Department of Microbiology with Parasitology and Immunology, Faculty of Medicine, University of Novi Sad, 21000 Novi Sad, Serbia; Department of Biology, Stanford University, Stanford, CA, USA; ChEM-H Institute, Stanford University, Stanford, CA 94305, USA; Chan Zuckerberg Biohub; San Francisco, CA 94158, USA

**Author notes:** These authors contributed equally (Tomás Cervantes Rincón, Tereza Frckova, Zaria I. Contejean). Equal senior contribution (Pavle Banović, Daniel Ruzek, Christopher O. Barnes, Davide F. Robbiani).

## Abstract

West Nile virus (WNV) is a mosquito-borne pathogen of global concern that can cause fatal neuroinvasive disease. No specific prophylaxis or treatment exists for WNV or related orthoflavivirus infections, and the determinants of human disease severity remain poorly understood. Here, we report that neutralizing autoantibodies against type I interferons do not impair antiviral antibody development. Among the fully human monoclonal antibodies with potent neutralizing activity against WNV that were discovered, W010 targets a unique epitope within the envelope protein domain III (EDIII) and confers both pre- and post-exposure protection in a murine WNV model, even when interferon signaling is impaired. A second protective antibody, W014, exhibits broad cross-neutralization of other pathogenic orthoflavivirus members, including Japanese encephalitis virus, Murray Valley encephalitis virus, Saint Louis encephalitis virus, and Usutu virus. These findings identify key neutralizing epitopes on WNV EDIII and provide candidates for the development of antibody-based interventions against encephalitic orthoflavivirus infections.

## INTRODUCTION

West Nile virus (WNV; *Orthoflavivirus nilense*) is one of many orthoflaviviruses (genus *Orthoflavivirus*) transmitted by *Culex* spp. mosquitoes that are responsible for emerging infectious diseases globally, including fatal encephalitis.^1-4^ While most human WNV infections are asymptomatic, an estimated 20 to 40 percent result in mild disease known as West Nile fever (WNF) and approximately 1 percent progress to neuroinvasive disease (WND), which can be lethal.^5-8^ The reasons for the difference in disease trajectory upon WNV infection are multifactorial; established risk factors include advanced age and CCR5 polymorphisms.^9-11^ In addition, emerging evidence indicates that the presence of autoantibodies neutralizing type I interferons (IFN-Is: IFN-α and/or IFN-ω) may also impact disease upon infection by WNV and other orthoflaviviruses.^8,12-16^

WNV is globally endemic and remains the primary cause of mosquito-borne illness in the United States. Since the large outbreak of 2018, it has also become an increasing public health concern in Europe.^17-20^ Projections indicate that the incidence of WND will continue to rise and that the virus may expand into new regions, likely driven by climate change and the associated spread of mosquito vectors.^21^ Despite representing a considerable public health threat, there are no specific prophylactic means or treatments available against WNV-related human diseases.^22^

Monoclonal antibodies have demonstrated clinical efficacy against a growing number of infectious diseases, particularly when administered early after the exposure or the onset of symptoms.^23-25^ Although no monoclonal antibody has yet been approved for the prevention or treatment of orthoflavivirus infections, several candidates against yellow fever, dengue, Zika, and tick-borne encephalitis are currently in clinical development.^26-28^ For WND, a humanized mouse-derived monoclonal antibody has progressed to clinical development.^29^ Of note, transfer of polyclonal immunoglobulins with relatively high titers of anti-WNV antibody failed to show efficacy in humans when administered during late stages of severe disease.^30^

Like other members of the *Orthoflavivirus* genus, WNV displays a single envelope (E) protein on its surface that mediates viral entry.^5,31^ The E protein comprises three domains (I, II, III), with E domain III (EDIII) forming an immunoglobulin-like fold that plays a key role in attachment to host cells.^4,32^ Antibodies against EDIII have been identified as potent neutralizers of various orthoflaviviruses, including those transmitted to humans by *Aedes* spp. mosquitoes and ticks.^32-39^ While monoclonal antibodies targeting the WNV EDIII domain have been previously described,^40,41^ these were either of mouse origin or derived from individuals not selected for high serum neutralizing activity. Although some antibodies showed neutralizing activity against WNV infection, they lacked cross-reactivity with serologically related orthoflaviviruses, such as Japanese encephalitis virus (JEV), Murray valley encephalitis virus (MVEV), Saint Louis encephalitis virus (SLEV), or Usutu virus (USUV), all of which are capable of causing severe, and sometimes fatal, human disease.^3,42,43^

Here, we report the discovery and characterization of human monoclonal antibodies that potently neutralize WNV, some of which also exhibit cross-neutralizing activity against other mosquito-borne orthoflaviviruses. We identify and characterize the epitope of 4 antibodies, including W010, a highly potent monoclonal antibody that confers both pre- and post-exposure protection against WNV in mice, and W014, which demonstrates broad neutralizing activity against related orthoflaviviruses.

## RESULTS

### Serum antiviral antibodies and autoantibodies in a West Nile disease cohort

Seventy-two study participants with suspected WND or WNF were enrolled during 2022 and 2023 in Serbia, and their serum collected during hospitalization (Figure S1A and B and Table S1). Serially diluted samples were tested by the enzyme-linked immunosorbent assay (ELISA) for the presence of IgG antibodies targeting the EDIII protein of WNV (WNV_EDIII_; Figure 1A). As expected, based on previous studies of the antibody response to other orthoflaviviruses,^34,44,45^ the amount of IgG antibodies to the WNV_EDIII_ was broadly variable.

**Figure 1.**
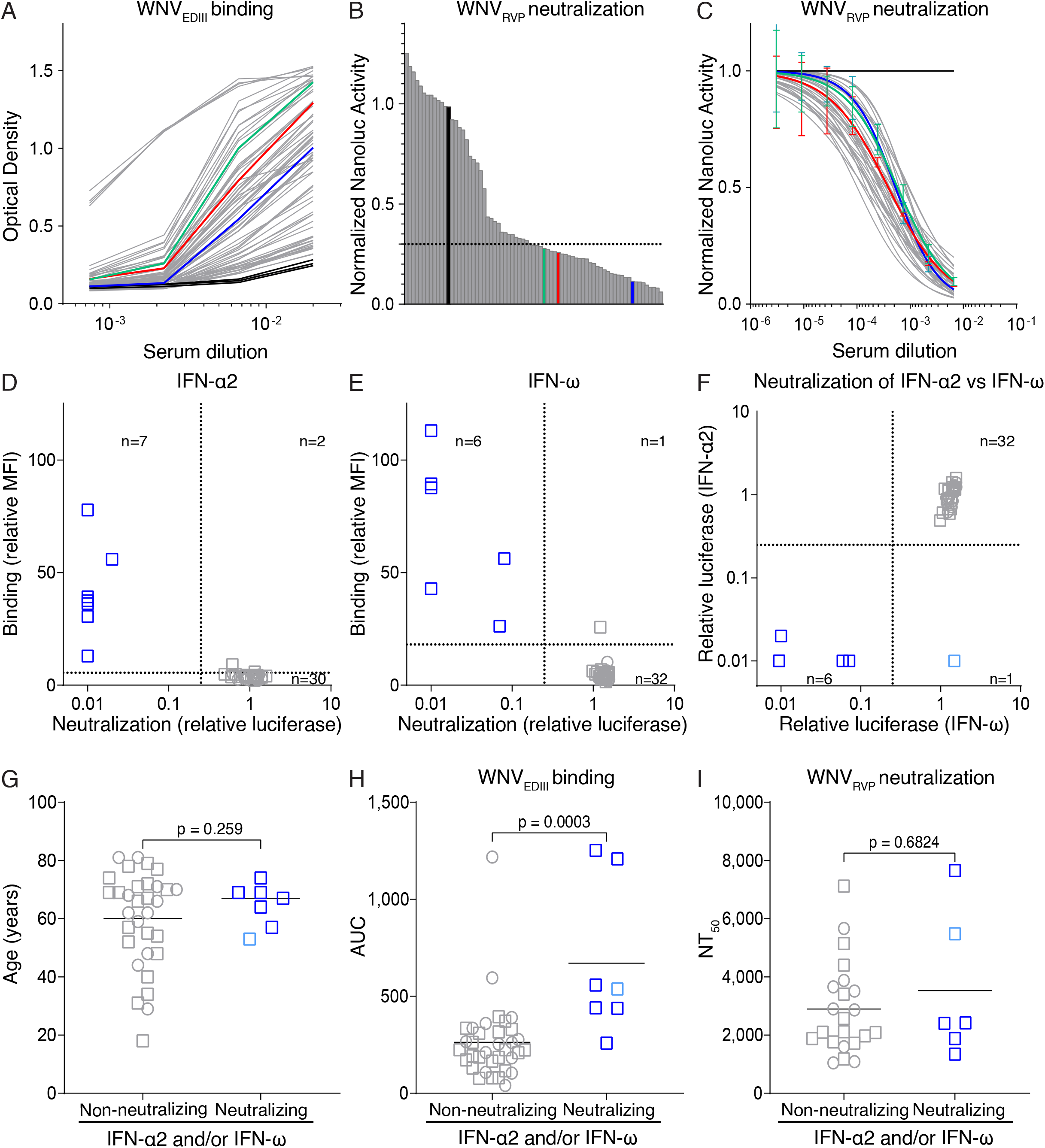
Serum antibodies and autoantibodies in a cohort of individuals hospitalized for WND/WNF. **(A)** IgG binding to WNV_EDIII_ by serial serum dilution. ELISA curves for samples from the 72 WND/WNF-suspected individuals and 3 orthoflavivirus naïve controls (black) are shown. The average of two independent experiments is shown. **(B)** Serum neutralization screening with WNV_RVP_. Shown is the rank-ordered NanoLuc activity relative to no-serum control for the 72 WND/WNF-suspected individuals and 1 orthoflavivirus naïve control (black); lower values correspond to higher neutralization. Samples with relative NanoLuc signal below 0.3 (dotted line) were selected for neutralization curves in (C). Each bar represents and individual participant sample analyzed at 1:100 dilution. Average signal of triplicate wells from a single experiment. **(C)** Neutralization of WNV_RVP_ by serial serum dilution. Shown is the NanoLuc activity relative to no-serum control for 36 WND/WNF cases and 1 orthoflavivirus naïve control (black). Mean ± SD of triplicates. Representative of two independent experiments. **(D-F)** Identification of serum autoantibodies to IFN-α2 and IFN-ω. Plots compare the ability of serum IgG to bind, and of serum to neutralize, IFN-α2 (D) and IFN-ω (E) (n=39; only samples for which sufficient serum was available were assayed). The comparison of IFN-α2 and IFN-ω neutralization is shown in (F). Binding is shown as relative Mean Fluorescence Intensity (MFI) and neutralization as relative ISG15-promoter driven luciferase signal compared to no serum control. The dotted lines indicate the threshold for positivity of the assay.^47,78^ Representative of 2 independent experiments. **(G)** Age and gender distribution of the study participants with or without IFN-α2 and/or IFN-ω neutralizing autoantibodies. Welch’s test. **(H and I)** Serum IgG binding to WNV_EDIII_ and serum neutralization of WNV_RVP_ in study participants with or without IFN-α2 and/or IFN-ω neutralizing autoantibodies. The two groups are compared with respect to (H) IgG binding to WNV_EDIII_ (Area Under the Curve (AUC) of ELISA and (I) neutralization of WNV_RVP_ (NT_50_ values). Horizontal lines indicate the mean. Mann-Whitney test. In (A to C), green, blue and red indicate samples from WNV-infected individuals from which antibodies were derived (Figure S2B and C). In (D to I), female is circle, male is square, dark blue neutralizes both IFN-α2 and IFN-ω, light blue neutralizes IFN-α2 only, grey does not neutralize either.

To assess the virus neutralizing activity of serum, samples were evaluated using an improved WNV reporter virus particle system with NanoLuc luciferase (WNV_RVP_, see Material and Methods; Figure S1C and 1B). Thirty-six individuals with high serum neutralizing activity in the screening were selected for analysis by serial serum dilution (Figure 1C). Half-maximal neutralizing titers (NT_50_) were variable, with no significant correlation between anti-WNV_EDIII_ IgG levels and WNV_RVP_ NT_50_ values (Figure S1D). Furthermore, no significant correlation or difference were noted in IgG levels to WNV_EDIII_ or WNV_RVP_ neutralization when these features were examined against demographic characteristics, days of hospitalization, or survival, with the exception of a weakly significant positive correlation between IgG levels to WNV_EDIII_ and age, and lower NT_50_ values for individuals who succumbed to the disease (Figure S1E to N).

Serum autoantibodies neutralizing IFN-α and/or IFN-ω were recently identified as a predisposing factor for WND.^8^ To establish their presence in the cohort and possible association with distinct WNV serologic features, we used a multiplexed bead-based assay to measure IgG autoantibodies binding to IFN-α2 or IFN-ω^46,47^ and a recently developed highly sensitive biological assay to determine IFN-α and/or IFN-ω neutralization.^48^ Ten out of 39 individuals with WND (26%) that were analysed showed detectable levels of serum IgG targeting IFN-α2 and/or IFN-ω, and seven of these reactive sera (18%) could neutralize low IFN-α2 (0.01 ng/mL) and/or low IFN-ω (0.02 ng/mL) doses (Figure 1D to F). These frequencies represent a significantly higher prevalence than in the general population, as shown by comparison to blood donors and in recent reports^46,49^ (Figure S1O and P). The individuals with neutralizing IFN-α and/or IFN-ω autoantibodies were all male and on average slightly older than individuals without these autoantibodies (64.7 vs 60 years) (Figure 1G). Notably, the presence of autoantibodies neutralizing IFN-α and/or IFN-ω did not adversely impact the serologic response to WNV. On the contrary, individuals with these autoantibodies exhibited a significantly increased IgG response to the WNV_EDIII_ protein, while their WNV_RVP_ NT_50_ values were comparable to those of individuals without such autoantibodies (Figure 1H and I). We conclude that the development of antiviral antibodies in individuals hospitalized due to WNV infection is not impaired by the presence of autoantibodies against IFN-α and/or IFN-ω.

### Features of human WNV_EDIII_ antibodies

To characterize anti-WNV antibodies, we isolated by fluorescence-activated cell sorting (FACS) WNV_EDIII_-specific B cells from the peripheral blood mononuclear cells (PBMCs) of three convalescent participants with high WNV_RVP_ neutralizing activity (NT_50_ values ranging between 1717 and 2567) and no detectable autoantibodies against IFN-α and/or IFN-ω in serum (Figure 1C, S2A and B). In total, 107 antibody heavy and light chain gene pairs (89 IgG, 14 IgM and 4 IgA) were amplified by RT-PCR from individually purified memory B cells and sequenced. Clonally expanded antibody sequences were found in all 3 participants (Figure S2C), and the CDR3 length, as well as the overall level of antibody V gene somatic hypermutation, were similar to previous reports (Figure S2D).^32,34,50^

Forty-four antibodies (including at least one representative for most of the expanded clones) were expressed recombinantly as IgG1 molecules. The half-maximal effective concentration (EC_50_) values for binding to the WNV_EDIII_ in ELISA was obtained for 32 of them (72.7%) and, consistent with the high similarity at the EDIII between lineage I and II of WNV (the two lineages responsible for most human disease),^5,19^ their binding to the corresponding EDIII was very similar (Figure 2A and B, S2E and Table S2). To determine whether, in addition to WNV, the monoclonal antibodies were reactive to the EDIII of related orthoflaviviruses of the JE serocomplex, we produced the EDIII proteins corresponding to JEV, MVEV, SLEV, and USUV for use in ELISA. Some of the antibodies displayed cross-reactivity, including W014 (JEV, MVEV, SLEV, and USUV), W011 (JEV, MVEV, and USUV), and W051 (MVEV and SLEV) (Figure 2A and Table S2). For most of the antibodies, no binding was noted to the EDIII of orthoflaviviruses transmitted by ticks (tick-borne encephalitis virus, Powassan virus, Kyasanur forest disease virus, Langat virus, Louping-ill virus (LIV), Omsk haemorrhagic fever virus) or *Aedes* spp. mosquitoes (four serotypes of dengue virus (DENV1-4), yellow fever virus, Zika virus (ZIKV)), with the exception of W006 (LIV), W023 (ZIKV), and W030 (DENV3, DENV4 and ZIKV) (data not shown). Thus, EDIII antibodies elicited by WNV infection can be either specific for WNV or react more broadly with related orthoflaviviruses.

**Figure 2.**
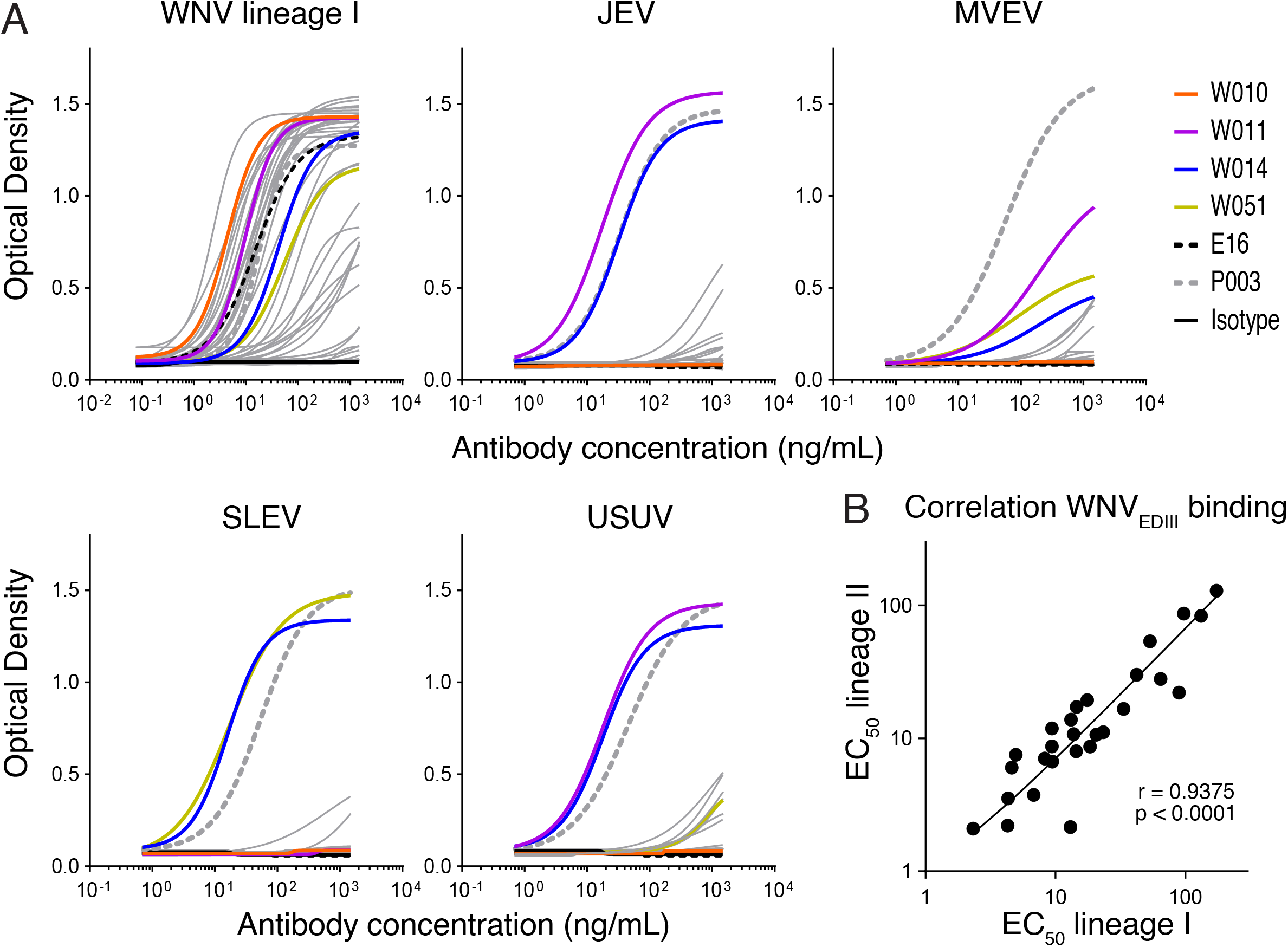
Human monoclonal antibodies isolation and crossreactivity. **(A)** Monoclonal antibody binding to the EDIII of WNV and related orthoflaviviruses. ELISA curves show the binding of 44 monoclonal antibodies to the EDIII of the indicated orthoflaviviruses transmitted by *Culex spp*. mosquitoes. Positive control is the broadly reactive antibody P003;^79^ humanized mouse antibody E16^41^ is shown for comparison. Representative of two independent experiments. **(B)** Correlation of the monoclonal antibodies binding to the EDIII corresponding to WNV lineages I and II. Each dot represents an individual monoclonal antibody EC_50_ value. Pearson correlation.

### Low dose W010 protects mice even five days after WNV challenge

When evaluated for neutralizing activity, 15 of the 44 antibodies (34%) showed robust WNV_RVP_ neutralization (half-maximal inhibitory concentration -IC_50_-values below 10μg/mL) and two outstanding antibodies were identified: W010 and W049 (IC_50_ values of 3.7 and 2.0 ng/mL, respectively), both of which outperformed the humanized mouse-derived antibody E16^29^ (Figure 3A and Table S2). Select antibodies were further tested against authentic WNV lineage I and II by two complementary virus neutralization assays, which confirmed the antibodies neutralizing capacity and identified W010 as the most potent against both lineages (IC_50_ values: 4.8 ng/mL for lineage I and 3.8 ng/mL for lineage II when measuring the cytopathic effect; Figures 3B and S3A).

**Figure 3.**
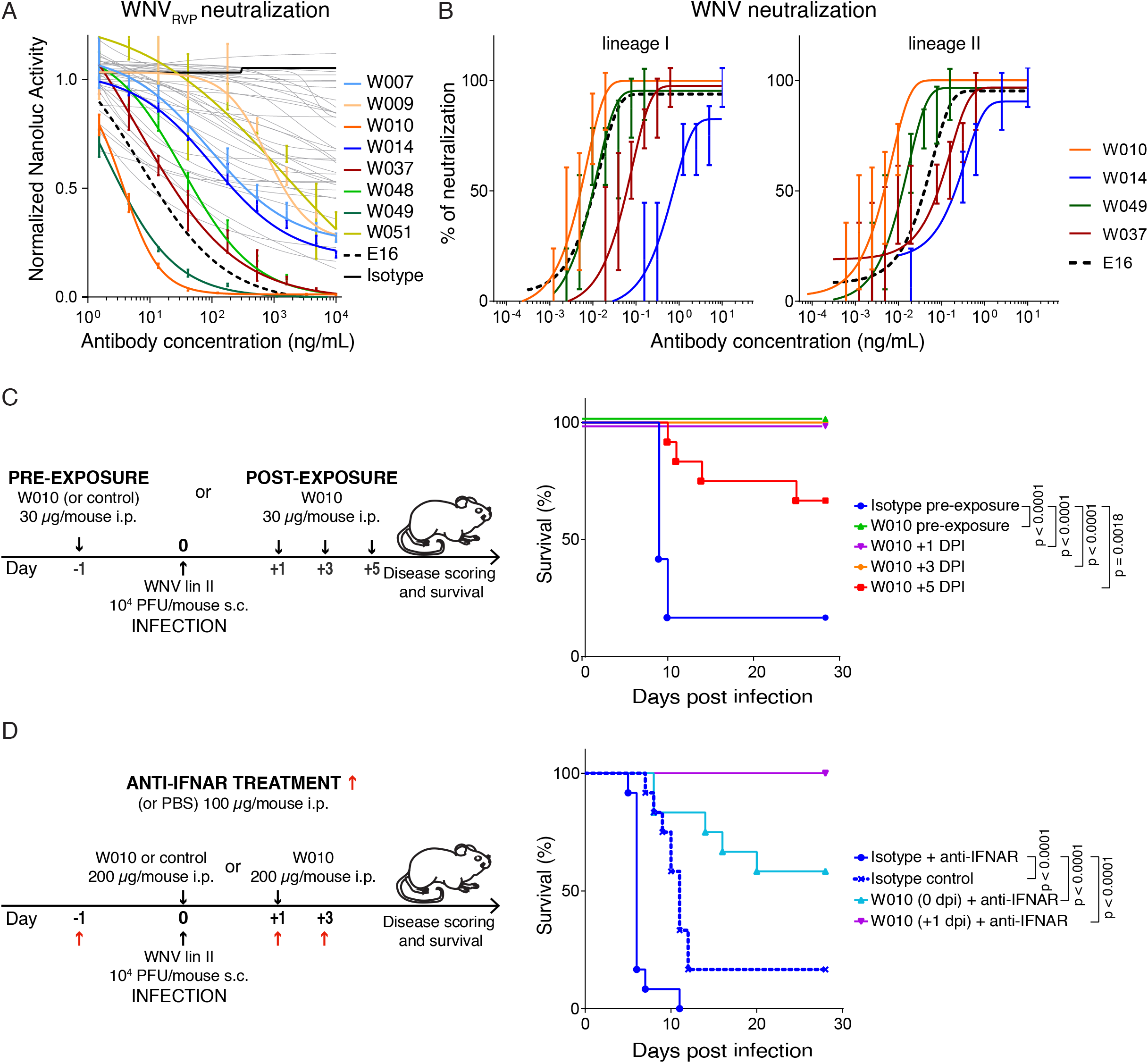
Identification of W010 as potent neutralizer of WNV. **(A)** Neutralization of WNV_RVP_ by monoclonal antibodies. NanoLuc activity in the presence of serially diluted antibodies relative to no-antibody control is shown. Mean ± SD of triplicates. Representative of two experiments. **(B)** Neutralization of virus isolates representing WNV lineages I and II. Representative neutralization curves by serially diluted antibodies. The percentage of neutralization was calculated based on the development of cytopathic effects 5 days after infection by comparing antibody-treated to non-treated control. Mean ± SD of octuplicates. Representative of two experiments. See also Figure S3A. **(C)** Efficacy of W010 in pre- and post-exposure prophylaxis against lethal WNV lineage II infection. Left, diagram of the experiment’s timeline. Right, survival curves of each experimental group are compared. Two independent experiments with six mice per group in each experiment were combined and differences in survival were analysed by log rank Mantel-Cox test. See also Figure S3B. **(D)** Therapeutic efficacy of W010 against lethal WNV lineage II infection of mice receiving anti-IFNAR antibodies. Left, diagram of the experiment’s timeline. Right, mouse survival over time. Two independent experiments with six mice per group in each experiment were combined and differences in survival were analysed by log rank Mantel-Cox test. See also Figure S3C.

To assess whether W010 protects against disease, we first conducted pre- and post-exposure prophylaxis experiments with BALB/c mice (Figure 3C and S3B). Ten of 12 control mice (83%) that received isotype antibody 24 hours prior to lethal challenge with 10^4^ plaque-forming units (PFU) of WNV lineage II succumbed to the infection by day 10. In contrast, all mice treated with a low dose of W010 (30 µg/mouse) at either 1 day before, or 1 or 3 days after infection, were protected from disease (p<0.0001 for each group compared to isotype control). Notably, low-dose W010 was also significantly protective when administered 5 days post infection (p=0.0018).

Next, to mimic the high-risk situation of individuals with autoantibodies to IFN-α and/or IFN-ω, we evaluated the protective effect of W010 in mice conditioned with antibodies blocking the IFN-I receptor (interferon alpha/beta receptor subunit 1, IFNAR; see Material and Methods; Figure 3D and S3C). When isotype antibody was administered on the day of WNV infection, all 12 mice succumbed by day 11. Remarkably, 58% and 100% of mice receiving W010 on the day of infection or 1 day post infection, respectively, survived (n=12 and p=0.0001 for both compared to isotype control). W010 is not polyreactive, displays an average half-life of 104 hours in FcγR/FcRn-humanized mice^51^ and shows a melting temperature of 79.6°C (Figure S3D to F). Altogether, these results demonstrate the protective efficacy of W010 even in the context of impaired IFN-I signaling, supporting its potential for prevention and therapy against WNV also in individuals harboring IFN-I autoantibodies.

### Anti-WNV neutralizing antibodies distinctly target the lateral ridge of EDIII

To approximate the epitope recognized by anti-WNV antibodies, we performed competition binding experiments using surface plasmon resonance (SPR). By this method, neutralizing antibodies could be classified into three main competition bins: W009, W010, W048 and W049 competed with one another, but not with W007 and W014, whereas W037 competed with all antibodies tested (Figure 4A and S4A and B). To define the structural basis underlying these competition profiles and associated neutralization mechanisms, we solved high-resolution crystal structures of the antigen-binding fragment (Fab) from W010, W014, W037, and W049 in complex with the WNV_EDIII_ protein. Consistent with the SPR competition data, the antibodies adopt distinct binding orientations allowing recognition of distinct epitopes on EDIII (Figure 4B and C).

**Figure 4.**
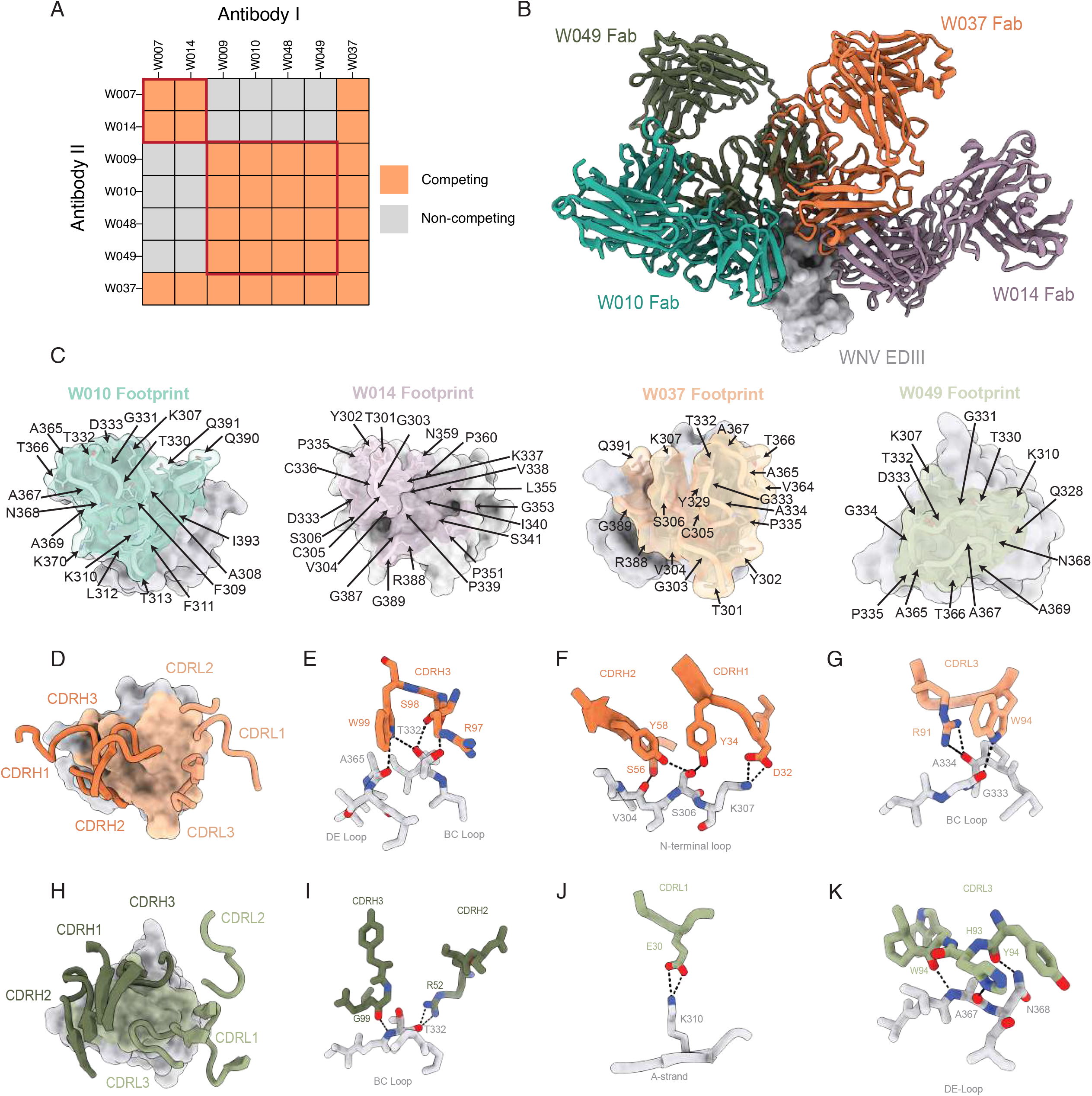
Characterization of the EDIII neutralizing epitopes. **(A)** Epitope binning by competition SPR. Upon immobilization of Antibody I, WNV_EDIII_ was bound, followed by Antibody II. Summary of antibodies competing for the same epitope by this method. See also Figure S7. Related to Figure S4. **(B)** Superposition of crystal structures of WNV_EDIII_ in complex with W010, W037, W014 and W049 Fabs resolved to 1.8Å, 2.1Å, 1.4Å and 2.5Å, respectively. Structures were aligned to the WNV_EDIII_. **(C)** The EDIII epitopes of W010, W014, W037 and W049 defined as interfacing residues in PDBePISA (see methods). **(D)** W037 Fab CDR loops mapped onto the antibody footprint, which is highlighted in orange on the EDIII surface. **(E)** Molecular interactions between W037 CDRH3 loop and the EDIII BC and DE loops. Trp99_HC_ mediates hydrogen bonding with backbone atoms Ala365_EDIII_ and Thr332_EDIII_. **(F)** W037 CDRH2 and CDRH1 residues form a series of potential hydrogen bonds with the N-terminal loop region of the lateral ridge epitope, including a common mode of engagement by the side chain of Tyr58_HC_ and Tyr34_HC_ with Ser306_EDIII_. **(G)** W037 CDRL3 engagement with the BC loop region of the lateral ridge epitope utilizes a combination of hydrophobic and hydrogen bonding interactions. **(H)** W049 Fab CDR loops mapped onto the antibody footprint, which is highlighted in green on the EDIII surface. **(I)** W049 CDRH3 and CDRH2 hydrogen bonding interactions between Gly99_CDRH3_ and Arg52_CDRH2_ with Thr332_EDIII_ in the BC loop. **(K)** W049 CDRL1 mediates salt bridge formation between Glu30_EDIII_ and Lys310_EDIII_ in the A-strand of the lateral ridge. **(L)** W049 CDRL3 utilized a combination of charged electrostatic and hydrogen-bonding interactions involving DE loop residues Ala367_EDIII_ and Asn368_EDIII_.

The structure of the W037 Fab-WNV_EDIII_ complex revealed recognition of the lateral ridge, encompassing the N-terminal region and the BC, DE, and FG loops of EDIII, which together span the EDI-EDIII interface^52^ (Figure 4D to G).

Similarly, W049 engages the lateral ridge epitope in a manner closely resembling that of the mouse anti-WNV antibody E16, with primary contacts involving the A-strand and the BC and DE loops (Figure 4H to K and Figure S4C). W049 binding is mediated through CDRH3 and CDRL3 contacts with the BC and DE loops, forming an extensive hydrogen-bonding network centered on Thr332_EDIII_ and Asn368_EDIII_. Additionally, the CDRL1 of W049 forms a stabilizing salt bridge between Glu30_CDRL1_ and Lys310_EDIII_. Together, W037 and W049 represent canonical lateral ridge binding antibodies, and share a binding orientation similar to potent monoclonal antibodies against several orthoflaviviruses (Figure S4C and D). In contrast, W010 and W014 exhibited unique binding poses not previously observed.

The structure of the W010 Fab-WNV_EDIII_ complex revealed engagement of the A-strand, and of the BC and DE loops, mediated primarily through the CDRH3 and CDRL3 loops (Figure 5). W010 buries a total surface area (BSA) of 705.5Å^2^ on EDIII, with contributions from both heavy (360Å^2^) and light (345.6Å^2^) chains. Key CDRH3-interactions include a salt bridge between Glu100_G-HC_ and Lys310_EDIII_ on the A-strand, as well as side-chain and backbone interactions with residues in the BC and DE loops by Glu100_E-HC_ and Cys100_B-HC_, respectively (Figure 5B and C). Lys310_EDIII_ is also engaged by CDRL3 residues Ser91_LC_ and Tyr92_LC_ (Figure 5D). Given that Lys310_EDIII_ accounts for ∼14% of the total BSA of the epitope, low sequence conservancy at this position among the orthoflaviviruses of the JE serocomplex likely explains W010’s lack of cross-reactivity (Figure 5E).

**Figure 5.**
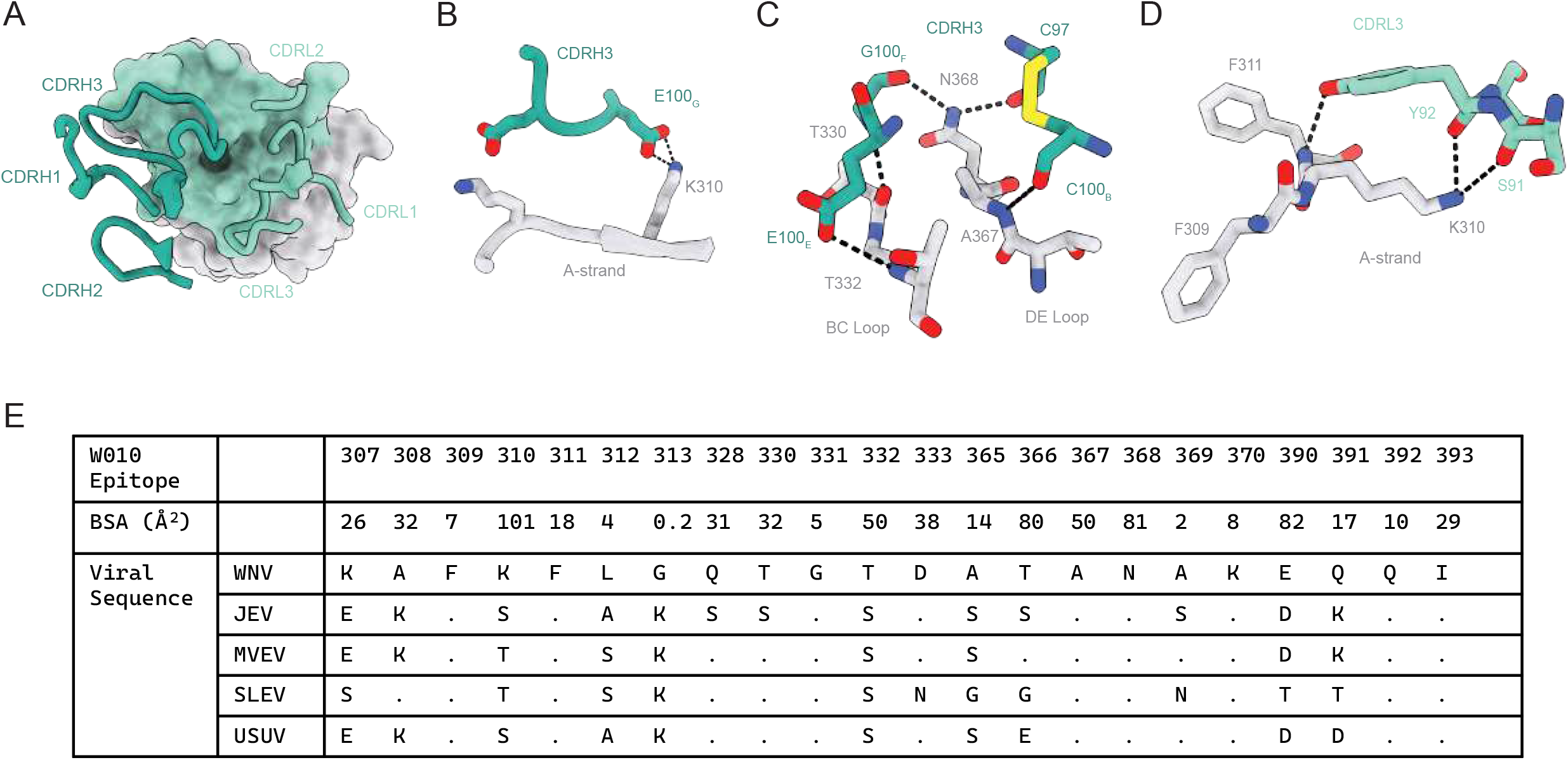
Characterization of the neutralizing epitope of W010. **(A)** W010 Fab CDR loops mapped onto the antibody footprint, which is highlighted in cyan on the EDIII surface. **(B)** W010 CDRH3 interactions with WNV_EDIII_ display salt bridge between residue Glu100_G-HC_ and Lys310_EDIII_ in the A-stand of the lateral ridge epitope. **(C)** Hydrogen bonding networks between W010 CDRH3 residues and the BC and DE loops of the EDIII lateral ridge epitope. **(D)** W010 CDRL3 interactions with WNV_EDIII_. Hydrogen bonds between Ser91-Tyr92_CDRL3_ and Lys310_EDIII_ are shown. Hydrogen-bonds are represented as dashed lines between donor and acceptor atoms. **(E)** Sequence contacts for W010 epitope and associated combined V_H_- and V_L_-buried surface area. The WNV EDIII is used as the reference sequence for alignment with EDIII proteins of the JE serocomplex. Identical residues in the JE serocomplex are indicated with a period and differences are represented by the residue at that position.

To gain insight about how W010 binds to and neutralizes the virus, we used a previously determined cryo-electron microscopy (cryo-EM) structure of the WNV virion to model epitope recognition^53^ (Figure S5A to C). In the context of the mature WNV virion, the W010 epitope is fully accessible, with W010 anchoring at the loop regions directly adjacent to the EDII domain of the same protomer that comprise the lateral ridge epitope. W010’s pose provides a binding advantage whereby two Fab molecules can engage the WNV epitope at the 5-fold symmetry axis suggesting a mechanism for enhanced potency through avidity effects (Figure S5 B). Consistent with this view, W010 IgG is superior to Fab in WNV neutralization (Figure S5D). Thus, both the mode of engagement and bivalent binding contribute to the potency of W010. Passaging the virus under subneutralizing concentrations of W010 *in vitro* led to the emergence of antibody-resistant variant carrying the amino acid change N368T, a residue contributing ∼11% of the epitope’s total BSA (Figure 5E and data not shown). While the structure reveals a direct hydrogen bond between N368 and the antibody, supporting its role in epitope recognition (Figure 5C), the structural data alone do not fully explain the magnitude of the ∼100-fold reduction in neutralization observed upon introduction of the N368T substitution (Figure S5E). However, the functional importance of this residue is supported by both the observed contact in the structure and the virus escape data.

### Identification of a protective epitope shared by the Japanese encephalitis serocomplex

WNV_EDIII_ antibodies were next screened for the ability to cross-neutralize RVPs corresponding to related viruses of the JE serocomplex (see Material and Methods). W014 (clonally related to W007) was identified as a broad neutralizer of WNV (IC_50_ value = 102 ng/mL), JEV (IC_50_ value = 43.6 ng/mL), MVEV (IC_50_ value = 27.7 ng/mL), SLEV (IC_50_ value = 4.3 ng/mL), and USUV (IC_50_ value = 36.2 ng/mL), while W051 neutralized MVEV (IC_50_ = 485 ng/mL) and SLEV (IC_50_ = 12.8 ng/mL) (Figure 6A and S6A, Table S2). In line with these observations, sera from the three individuals, from whom antibodies were derived, also exhibited cross-neutralization of JEV, MVEV, SLEV, and USUV RVPs (Figure S6B).

**Figure 6.**
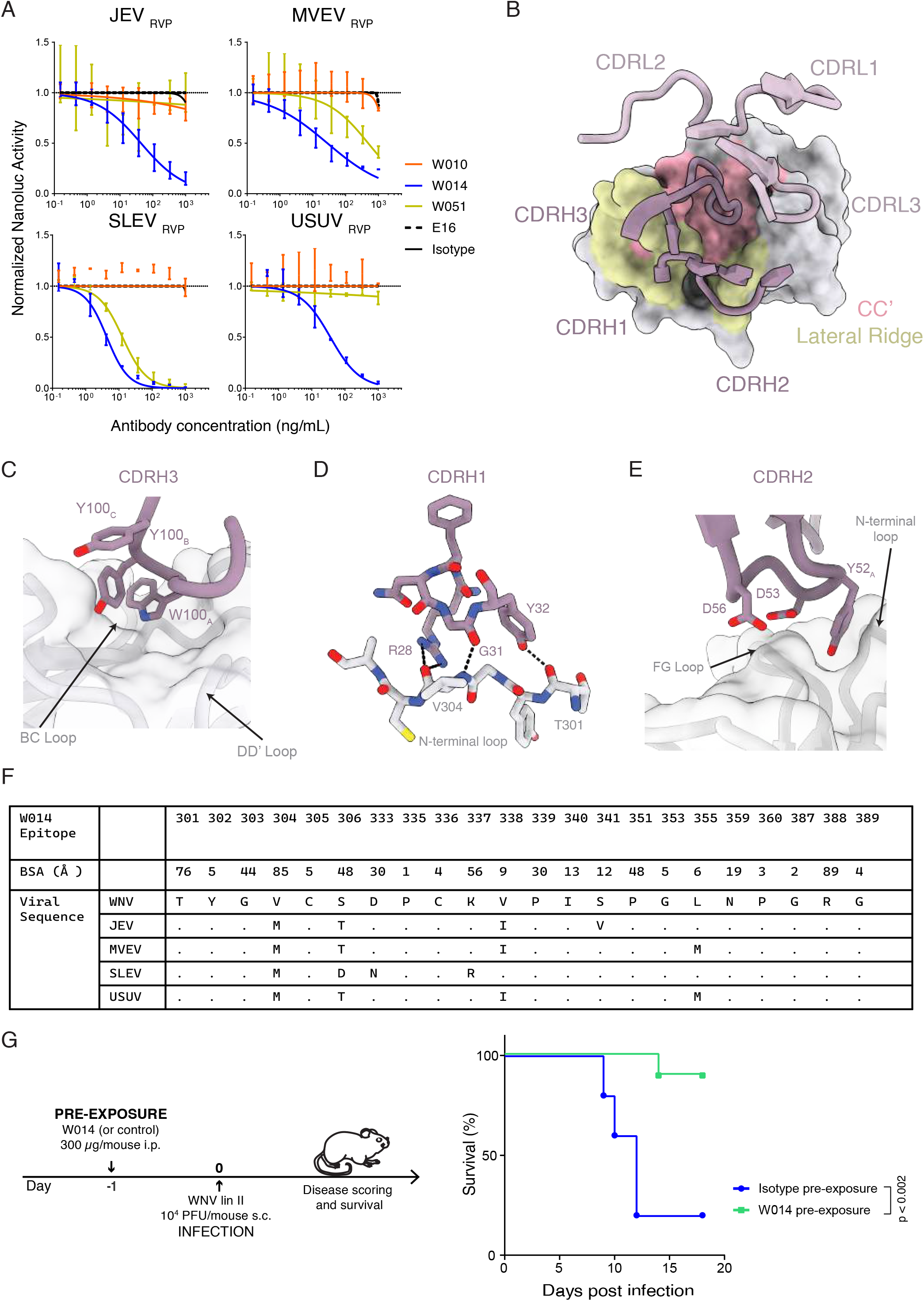
Structural basis of the broadly neutralizing epitope of W014. **(A)** Neutralization of RVPs corresponding to WNV-related orthoflaviviruses JEV, MVEV, SLEV and USUV. NanoLuc activity in the presence of serially diluted antibodies relative to no-antibody control is shown. Mean ± SD of triplicates. Representative of two experiments. **(B)** W014 CDR loops interface with WNV EDIII lateral ridge (yellow) and CC’ (pink) epitopes. **(C)** Surface and cartoon representation of the CC’ hydrophobic pocket recognized by the aromatic residues at the tip of the W014 CDRH3 loop. **(D)** W014 CDRH1, primarily uses hydrogen-bonding contacts with both side-chain and backbone peptide atoms to interface with the N-terminal loop. **(E)** W014 CDRH2 buries Tyr52_A_ into a hydrophobic cleft formed by the N-terminal and the FG loops of the lateral ridge epitope. **(F)** Sequence contacts for W014 epitope and associated combined V_H_- and V_L_-buried surface area. The WNV EDIII is used as the reference sequence for alignment with EDIII proteins of the JE serocomplex. Identical residues in the JE serocomplex are indicated with a period and differences are represented by the residue at that position. **(G)** Efficacy of W014 in pre-exposure prophylaxis against lethal WNV lineage II infection. Left, diagram of the experiment’s timeline. Right, survival curves of each experimental group are compared. One experiment with ten (W014) and five (isotype) mice per group. Log rank Mantel-Cox test.

Structural analyses of the W014 Fab-WNV EDIII complex revealed recognition of the N-terminal and FG loops, which are canonical components of the lateral ridge epitope, and the CC’ and DD’ loops associated with the CC’ epitope (Figure 6B to E). In contrast to W010, the W014 light chain makes minimal contacts at the epitope, where interactions are mediated by CDRH1, CDRH2 and CDRH3 loops (V_H_ : 566Å^2^, V_L_ : 27.9Å^2^). W014 inserts an aromatic motif at the tip of the CDRH3 (Trp100_A-HC_ -Tyr100_C-HC_) into a cleft framed by Lys337_EDIII_, Pro339_EDIII_, and Pro351_EDIII_, which are highly conserved residues in the JE serocomplex (Figure 6C and F). Additionally, W014 forms backbone interactions at the lateral ridge whereby CDRH1 residues Arg28_HC_ and Tyr32_HC_ hydrogen bond with the carbonyls of Val304_EDIII_ and Thr301_EDIII_, respectively, and CDRH2 residue Tyr52_HC_ inserts into a pocket formed by the FG loop and N-terminal loop (Figure 6D and E). Collectively, engagement with strictly conserved EDIII residues and extensive backbone interactions in the W014 epitope likely facilitate cross-reactive binding among the orthoflaviviruses of the JE serocomplex (Figure 6F). However, this cryptic epitope is buried in the context of the mature virions, likely explaining W014’s weaker neutralization potency relative to W010 (Figure S5C). Thus, in addition to potent but monospecific antibodies like W010, infection by WNV can induce antibodies that are broadly effective against other members of the JE serocomplex.

In agreement with the RVP neutralization data, W014 neutralizes authentic WNV lineages I and II, (IC_50_ values of 773 ng/mL and 143 ng/mL, respectively, as determined by cytopathic effect assays; Figure 3B), as well as WNV, JEV, MVEV and SLEV by an immunofluorescence-based assay, albeit requiring higher antibody concentrations (Figure S3A and S6C). Finally, to assess the efficacy of W014 *in vivo*, we infected mice with WNV one day after antibody administration. W014 significantly prevented mice from dying (p=0.002; Figure 6G and S6D), demonstrating that the conserved epitope targeted by W014 mediates protection in a preclinical model.

## DISCUSSION

Infection with mosquito-borne WNV can result in severe neurological complications (i.e., encephalitis, meningitis, flaccid paralysis) with potentially fatal outcomes. In the absence of specific treatments, WNV remains a public health concern, especially due to its continuous geographic expansion.^21,54-56^ Herein, we describe the identification of potent human neutralizing antibodies with therapeutic potential for prevention or treatment of WNV-associated disease. Notably, some of these antibodies also exhibit cross-neutralizing activity against related orthoflaviruses in the Japanese encephalitis serocomplex, including JEV, MVEV, SLEV, and USUV, all of which are a medically relevant source of severe human diseases. Moreover, we report that the presence of autoantibodies neutralizing IFN-Is does not impair the development of antiviral antibody responses.

Recent studies have associated the presence of autoantibodies neutralizing IFN-Is, particularly IFN-α2 and/or -ω, with increased risk of severe disease and fatal outcome following orthoflavivirus infection, as well as adverse events following vaccination against yellow fever.^13-15^ In cases of WNV encephalitis, around 40% of poor clinical outcomes were linked to IFN-α2 and/or IFN-ω neutralizing autoantibodies^8^, a frequency comparable to that observed in the cohort reported herein (18%). These findings underscore the key role of IFN-Is in antiviral defenses^57-59^ and are consistent with observations that mice genetically defective in IFN-signaling,^16,60^ such as *IFN-α/βR*^−/−,61^ *Stat1*^-/-,16^ and *IRF-7*^-/-,62^ are more susceptible to neuroinvasive disease upon WNV infection. Importantly, we find that antiviral antibodies are not impaired by the presence of IFN-α2 and/or IFN-ω neutralizing autoantibodies, which is consistent with reports that IFN-α2 and/or IFN-ω autoantibodies do not interfere with coronavirus disease 2019 (COVID-19) vaccinal responses.^63-65^ In fact, individuals with IFN-α2 and/or IFN-ω neutralizing autoantibodies in our study exhibited significantly higher levels of WNV EDIII-specific antibodies, potentially reflecting higher viremia due to functional impairment of IFN-α2 and/or IFN-ω signaling that drives a more robust humoral response. This hypothesis could be tested in future studies by comparing viral and antigenic loads in IFN-α2 and/or IFN-ω competent patients versus those that are functionally impaired. Notably, however, higher antiviral antibody levels did not translate into improved virus neutralizing activity by the serum.

Human antibodies specific for the EDIII of other orthoflaviviruses were previously shown to possess strong virus neutralizing capabilities and potential for clinical development. For example, T025, a TBEV_EDIII_ antibody, is protective even at low doses in murine models of tick-borne encephalitis,^34^ while ZIKV_EDIII_ monoclonal antibodies Z004 and Z021 show efficacy in mouse as well as in macaque models of Zika congenital syndrome.^32,66^ These anti-EDIII antibodies, whose efficacy is restricted to a single or only few orthoflaviviruses, share some structural features including the usage of the lateral ridge epitope at the hinge of the EDI-EDIII regions like observed for W037 (Figure S4D).

W010 and W049, the two most potent WNV-neutralizing antibodies from this study, share key epitope contact features, including salt bridge formation with Lys310_EDIII_ and hydrogen bonding networks involving Thr332_EDIII_ and Asn368_EDIII_. Despite these similar contact points, W010 uses a divergent binding pose to engage components of the lateral ridge epitope, which may contribute to its enhanced neutralization potency. To further explore this possibility, we compared W010 to the WNV_EDIII_ antibody of mouse origin (E16) under clinical development, which recognizes the canonical lateral ridge epitope but has decreased neutralization potency^41,67^ (Figures 3B and S4D). In orthoflaviviruses, effective neutralization typically requires engagement of approximately 30 antibody molecules targeting the lateral ridge of EDIII based on the virion’s size and icosahedral symmetry.^68^ Both E16 and W010 bind this accessible epitope and can exceed the neutralization threshold, with E16 reported to engage up to 120 binding sites.^68^ Our new data indicate that both antibodies are capable of bivalent binding, suggesting that differences in avidity alone do not fully account for the increased neutralization potency of W010 (Figure S5D). Instead, we speculate that the distinct binding orientation of W010 may allow more favorable engagement of EDIII across both mature and immature virions, potentially increasing epitope accessibility and functional occupancy at critical sites such as the 5-fold symmetry axis. Together, these features are likely contributors to the enhanced neutralization potency observed for W010 relative to E16.

In contrast to EDIII antibodies that are potent but monospecific, some EDIII antibodies were also discovered that possess broader spectrum efficacy. W014, for example, neutralizes RVPs (and to a lower efficacy authentic viruses) corresponding to nearly all orthoflaviviruses that are transmitted by *Culex* spp. mosquitoes and relevant for human disease (WNV, JEV, MVEV, SLEV, USUV). Similar broadly acting antibodies were identified that are effective against the tick-transmitted pathogenic orthoflavivruses TBEV, OHFV, and KFDV (T056 and T058)^34^ or those transmitted by *Aedes* spp. mosquitoes ZIKV and the four serotypes of DENV (Z039^32^ and unpublished data). Structural analyses aimed at elucidating the molecular basis of cross-reactivity by broadly neutralizing anti-EDIII antibodies revealed that these antibodies recognize the lateral ridge and CC⍰ epitopes. Notably, W014 adopts a binding orientation that enables interaction with highly conserved residues within the CC⍰ epitope, which is shared across members of the Japanese encephalitis serocomplex.^69^ However, this conserved epitope is not exposed on either mature or immature virions, suggesting that conformational rearrangements of the virus envelope are required to transiently expose the CC⍰ site. A similar binding mode was previously observed for the anti-DENV antibody E111, which supports the hypothesis that CC⍰-targeting antibodies neutralize infection by recognizing conformational intermediates that arise as the envelope protein transitions from homodimeric to homotrimeric states during membrane fusion.^70^ Collectively, the antibodies described here reveal two principal epitope-targeting strategies that can elicit potent and, in some cases, broadly neutralizing responses. These findings also provide a structural and mechanistic framework to inform the design of broad-spectrum ‘prototype’ immunotherapies and vaccines, with the potential to protect against both currently circulating orthoflaviviruses and those that may emerge in the future that pose a threat to human health.

The *in vivo* experiments highlight the exceptional efficacy of W010, even when administered prophylactically at low dose (30 µg/mouse) or therapeutically several days after lethal WNV challenge of mice. A higher dose (200 µg/mouse) is also protective in mice with impaired IFN signaling. In humans, WNV viremia is most often accompanied by mild symptoms, which are reported to begin after an incubation period of 2-14 days.^5,71,72^ It is conceivable that administration of W010 at this time could reduce viral loads, thus limiting the risk of progression to the neurologic phase of the disease. It is also possible that prophylactic administration of antibodies in the context of WNF/WND seasonal outbreaks or in response to early warning^73,74^ would benefit especially at-risk populations, such as older adults, immunocompromised individuals, and those with functionally impaired type I IFN signaling. Similarly, W014 protects mice against WNV and future studies will assess whether its administration could lessen the risk of disease by JEV and other orthoflaviviruses of the JE serocomplex.^75-77^

In summary, the identification of human antibodies that are potent and broadly active against WNV and related orthoflaviviruses provides new knowledge relevant to the development of medical countermeasures against severe diseases that lack specific treatments and represent an important unmet medical need globally.

## ACKNOWLEDGEMENTS

We thank the study participants and their families, as well as the medical personnel at the clinical centers that were involved. At the IRB, we are grateful to Valentina Cecchinato and Mariagrazia Uguccioni (Human Subject Research), Andrea Celoria (technical support), and Melissa Robbiani (critical review of the manuscript). At the Rockefeller University, we thank Michel C. Nussenzweig for providing unpublished reagents and Tyler Lewy for providing guidance on experiments with mice treated with anti-IFNAR. At the California Institute of Technology, we thank Jennifer R. Keeffe for providing unpublished reagents and for comments on the manuscript. At the Pasteur Institute Novi Sad, we are grateful to Verica Simin and Ivana Bogdan for technical support during sample collection. We also thank M. Dvorakova (Biology Centre of the Czech Academy of Sciences) for technical assistance, and M. Motta, M. Borri and S. Fontana (Servizio Trasfusionale della Svizzera italiana) for providing control serum samples. X-ray crystallographic data were collected at the Stanford Synchrotron Radiation Lightsource, SLAC National Accelerator Laboratory, which is supported by the U.S. Department of Energy, Office of Science, Office of Basic Energy Sciences under Contract No. DE-AC02-76SF00515. The SSRL Structural Molecular Biology Program is supported by the DOE Office of Biological and Environmental Research, and by the National Institutes of Health, National Institute of General Medical Sciences (P30GM133894). We thank Daniel Fernandez and Beamline 12-2 staff for their support. The work was supported by Swiss-Czech collaborative projects funded jointly by the Swiss National Science Foundation (IZSEZ0_217585 and 320030L_231465 to D.F.R.) and the Czech Science Foundation (25-18189L to D.R.), by the project ‘InFlaMe’ funded by the European Union under Grant Agreement No. 101191725 (to D.R.), by the National Institute of Virology and Bacteriology, Programme EXCELES, project no. LX22NPO5103 funded by the European Union-Next Generation EU (to D.R.), by the Strategy of the Czech Academy of Sciences, AV21-Virology and Antiviral Therapy (to D.R.) by project reg. no. CZ.02.01.01/00/23_021/0012621 (SPOPROVIR), co-funded by the European Union, by the Swiss National Science Foundation (320030_232029 to B.G.H.; 310030_215559 and 310030E_205323 to L.V.), and in part by the Swiss Vaccine Research Institute (SVRI), by NIH grants U01 AI151698 (United World Antiviral Research Network, UWARN), P01 AI138938, and U19 AI111825 (to D.F.R.), R01 AI124690 (M.R.M.), and R01AI137276 (to S.B.). The content is solely the responsibility of the authors and does not necessarily represent the official views of the NIH. This study was also possible thanks to the IRB-Rockefeller University partnership for infectious disease research, supported in part by a grant to the IRB from the Fondazione Leonardo, to the Balkan Association for Vector-Borne Diseases (www.bavbd.org), to the European Virus Archive GLOBAL (EVA-GLOBAL) project that has received funding from the European Union’s Horizon 2020 research and Innovation program under grant agreement No 871029, and in part to funds from the Chan Zuckerberg Biohub (C.O.B.) and the Howard Hughes Medical Institute Emerging Pathogens Initiative (C.O.B.). Additionally, C.O.B. is supported by the Howard Hughes Medical Institute Hanna Gray Fellowship, Rita Allen Foundation, Pew Biomedical Scholars Program, and is a Chan Zuckerberg Biohub investigator. Z.I.C. is supported by a Ford Foundation Fellowship.

## AUTHOR CONTRIBUTIONS

D.F.R. and D.R. conceptualized the study, T. Cervantes Rincon, T. Frckova, Z. I. Contejean, K.G., B.C., F.B., L.S., S.T., V.H., S.B., L.V., M.P., B.G.H., P.B., D.R., C.O.B., and D.F.R., designed and analyzed the experiments. P.B, J.G., D.M., M.V., S.S., and D.L. designed human subject research protocols. T. Cervantes Rincon, T. Frckova, Z. I. Contejean, J.C., K.G., B.C., F.B. L.S., D.J., S.T., R.K., A.R.E.R., M.C., E.G.N., J.M., and Z.K. produced reagents and carried out experiments. M.R.M. provided unpublished reagents. T. Cervantes Rincon, R.T., and T.K. provided reagents. Z.B-O., J.G., M.V., and Z.L. collected and processed blood samples. D.M., J.G., Z.B., M.V., Z.L., D.L., S.S., M.B., L.P-D., B.P.J., and J.G. recruited participants and executed human subject research protocols. T. Cervantes Rincon, J.C., J.G., D.M., Z.B-O, M.V., Z.L, D.L., S.S., M.B., L.P-D., B.P-J. and J.G. processed human samples. Z.I. Contejean and A.R.E.R. performed crystallography experiments. Z.I. Contejean and C.O.B. performed structural analysis. T. Frckova, M.C., J.H., J.S., V.H., and D.R. performed in vivo experiments. S.G.M. performed bioinformatic analysis. T. Cervantes Rincon, T.F., Z.I.C., J.C., D.R., C.O.B., and D.F.R. wrote the manuscript with input from all coauthors.

## SUPPLEMENTAL INFORMATION

**Document S1. Figures S1-S6**

**Table S1**. Characteristics of study participants.

**Table S2**. Effective and inhibitory concentrations of recombinantly expressed monoclonal antibodies. **Table S3**. Crystallographic data processing and refinement statistics.

## SUPPLEMENTARY TITLES AND FIGURE LEGENDS

**Figure S1. Demographics and serologic features of individuals hospitalized for WND/WNF. Related to main Figure 1**.

**(A)** Demographic characteristics of the cohort.

**(B)** Geographic distribution of the patients’ residency. Case distribution shows the involvement of 11 Districts, with clusters in Central Serbia in 2022 and Northern Serbia in 2023. Blue dots indicate 2022 cases (with a dominant cluster near the city of Kragujevac, Šumadija District), while red dots represent the residency of the 2023 cases (with a dominant cluster near the city of Novi Sad). The map was generated with QGIS v3.12 (QGIS Development Team, 2020) starting from the GADM database (v4.1, July 2022, https://gadm.org/).

**(C)** Comparison between NanoLuc and Renilla reporter virus particles (RVPs). WNV_RVP_ were produced side-by-side by transfecting packaging cells with a NanoLuc- or Renilla-expressing WNV replicon (see Materials and Methods). Different volumes of supernatant-containing RVPs harvested at 24h, 48h and 72h were added to Huh 7.5 cells and the luminescence measured from cell lysates. The lower background with NanoLuc results in improved signal-to-noise ratio. Dotted horizontal lines indicate the average luminescence that was measured in the absence of WNV_RVP_.

**(D)** Correlation between serum IgG binding to WNV_EDIII_ and NT_50_ values. AUC is Area Under the Curve of ELISA binding. Pearson correlation.

**(E)** Correlation between serum IgG binding to WNV_EDIII_ and age at hospitalization. Pearson correlation.

**(F)** Correlation between serum IgG binding to WNV_EDIII_ and time from symptoms onset to serum sampling. Pearson correlation.

**(G)** Comparison of serum IgG binding to WNV_EDIII_ between samples from male and female participants. Unpaired two-tailed t-test.

**(H)** Correlation between serum IgG binding to WNV_EDIII_ and days of hospitalization. Pearson correlation.

**(I)** Comparison of serum IgG binding to WNV_EDIII_ between samples from survivors and non-survivors. Unpaired two-tailed t-test.

**(J)** Correlation between WNV_RVP_ NT_50_ values and age at hospitalization. Pearson correlation.

**(K)** Correlation between WNV_RVP_ NT_50_ values and time from symptoms onset to serum sampling. Pearson correlation.

**(L)** Comparison of serum WNV_RVP_ NT_50_ values between samples from male and female participants. Unpaired two-tailed t-test.

**(M)** Correlation between serum WNV_RVP_ NT_50_ values and days of hospitalization. Pearson correlation.

**(N)** Comparison of WNV_RVP_ NT_50_ values between samples from survivors and non-survivors. Unpaired two-tailed t-test.

**(O)** Prevalence of autoantibodies against type I IFNs in the WND cohort compared with available general population data.^49^ One-sided Fisher’s exact test.

**(P)** Prevalence of autoantibodies against IFN-α2 in the WND cohort compared with blood donor control samples from Serbia (left) and Switzerland (right), as determined by ELISA. AUC is Area Under the Curve. The threshold for positivity (dotted line) was determined set at 5 Standard Deviations of the signal of the controls. One-sided Fisher’s exact test.

**Figure S2. Isolation of WNV**_**EDIII**_**-specific memory B cells and neutralization of WNV. Related to main Figure 2**.

**(A)** Gating strategy. Forward and side scatter (FSC and SCC, respectively) were used to gate on lymphocytes. Dump channel included CD3, CD8, CD14, CD16 and a viability dye. CD20^+^ B cells binding to the WNV_EDIII_ coupled with PE and AF647 were isolated (see Methods for details).

**(B)** Identification of WNVEDIII-specific B cells. Flow cytometry plots show WNVEDIII-specific B cells (gate) in three WND convalescents. Percentages refer to the gated double-positive cells.

**(C)** Analysis of antibody sequences. Pie charts show the distribution of antibody sequences from sorted B cells (b). Coloured slices indicate clonally related sequences, while singlets are in white. In the center of the pie is the total number of sequences derived from each convalescent individual.

**(D)** Number of antibody V gene somatic nucleotide mutations and amino acid length of the complementarity-determining region 3 (CDR3).

**(E)** Monoclonal antibody binding to the EDIII of WNV lineage II. ELISA curves show the binding of 44 monoclonal antibodies to the EDIII of protein of WNV lineage II. Positive control is the broadly reactive antibody P003;^79^ humanized mouse antibody E16^41^ is shown for comparison. Representative of two independent experiments.

**Figure S3. Efficacy of W010 and suitability for clinical development. Related to main Figure 3**.

**(A)** Representative qualitative immunofluorescence microscopy images are shown for neutralization of WNV lineage I and II by the indicated monoclonal antibodies. Green is viral antigen, and blue is cell nuclei (4’,6-diamidino-2-phenylindole (DAPI)). Scale bar indicates 500 µm.

**(B)** Mouse symptoms over time after WNV challenge. DPI is days post infection. Related to Figure 3C. (C) Symptoms over time after WNV challenge in mice treated with anti-IFNAR. Related to Figure 3D.

**(C)** Antibodies’ polyreactivity. Monoclonal antibodies binding to insulin, keyhole limpet hemocyanin (KLH), lipopolysaccharyd (LPS), single-stranded DNA (ssDNA) and double-stranded DNA (dsDNA) was evaluated by ELISA. ED38 is a polyreactive monoclonal antibody used as positive control.^80^

**(D)** Half-life of W010 and W014 in humanized mice^51^ upon intravenous injection (50µg).

**(E)** Melting temperature values of WNV antibodies.

**Figure S4. Kinetics of monoclonal antibodies binding to WNV**_**EDIII**_ **and epitope binning by SPR. Related to main Figure 4A**.

**(A)** Representative Surface Plasmon Resonance plots of the assay to determine the antibody’s binding to the EDIII antigen (KD values).

**(B)** Competition of binding experiments identify antibodies binding to non-overlapping epitopes on the EDIII of WNV. Dotted black lines indicate EDIII binding to the immobilized antibody, while dotted red lines show the association of the non-competing antibody (or lack thereof when the lines are absent).

**(C)** Superposition of crystal structures of WNV_EDIII_ in complex with W049 and E16 (PDB 1ZTX).

**(D)** Superposition of crystal structures of WNV_EDIII_ in complex with W010, W037, W014, and E16 Fabs and canonical lateral ridge antibodies against Zika virus (Z004 Fab, PDB 6UTA) and tick-borne encephalitis virus (T025 Fab, PDB 7LSE).

**Figure S5. W010 and W014 engage distinct epitopes. Related to main Figure 5**.

**(A)** W010-EDIII and W014-EDIII were aligned with mature WNV virion (PDB 7KVA), with associated RMSD values of 0.52 and 0.55 respectively.

**(B and C)** Modeling of W010 (B) and W014 (C) Fab fragments binding to their respective EDIII epitopes at the 5-fold symmetry axis of the WNV.

**(D)** Neutralization of WNV (either RVPs, top, or authentic virus, bottom) by W010, comparing IgG1 to either Fab or to one-armed IgG1 (one arm W010 or E16, the other isotype), as indicated. MNR is Molar Neutralization Ratio.^81^

**(E)** Neutralization of wild type (WT) and mutant N368T WNV_RVP_ by W010 and E16. NanoLuc activity in the presence of serially diluted antibodies relative to no-antibody control is shown. Mean ± SD of triplicates. Representative of two experiments.

**Figure S6. Cross-neutralization by monoclonal antibodies and sera. Related to main Figure 6**.

**(A)** Monoclonal antibodies were analysed at a single dilution (500 ng/mL) for the ability to neutralize RVPs corresponding to WNV, JEV, MVEV, SLEV and USUV. NanoLuc activity relative to “no-antibody” control is shown. Orange represents the antibodies for which the NanoLuc activity of the RVPs was quenched by more than 55%. A single experiment, average of triplicate wells. RLS is relative luciferase signal over no-antibody control.

**(B)** Sera cross-neutralization. Sera from WND convalescents, from which monoclonal antibodies were derived, cross-neutralize RVPs corresponding to members of the JE serocomplex, TBEV and ZIKV. Neutralization curves are shown on top, heatmap with NT_50_ values is at the bottom.

**(C)** Representative qualitative immunofluorescence microscopy images are shown for neutralization of JEV, MVEV and SLEV by W014 and isotype control. Concentrations are expressed in µg/mL. Green is viral antigen, and blue is cell nuclei (DAPI). Scale bar indicates 500 µm.

**(D)** Mouse symptoms over time after WNV challenge in the presence of W014 or isotype control. DPI is days post infection. Related to Figure 6G.

## Methods

### Ethical statement

The study was approved by the ethical committees of University Clinical Centre of Vojvodina (approval No. 00–59), Blood Transfusion Institute of Vojvodina (approval No. 02-01/23), University Clinical Centre of Kragujevac (approval No. 01/22-389), and Pasteur Institute Novi Sad (approval No. 10-49/1). The research was conducted in compliance with the principles outlined in the Declaration of Helsinki and adhered to the Patient Rights Law of the Republic of Serbia. Healthy, orthoflavivirus naïve samples were obtained in Switzerland (Comitato Etico del Canton Ticino: CE-3428).

### Patient recruitment

Eligible patients presented at the Clinics for Infectious Diseases of Niš, Novi Sad or Kragujevac during 2022 and 2023 with signs, symptoms and laboratory findings consistent with WNF and/or WND. All cases of WNF/WND were considered probable if the patient met any of the following clinical criteria: (i) clinical presentation of encephalitis and/or meningitis and/or fever ≥38°C with (ii) presence of anti-WNV IgG in serum with seroconversion or a 4-fold increase in IgG titre on 2 subsequent samples (Euroimmun, Lubeck, Germany; EI 2662-9601 G). Demographic and clinical information on the cohort can be found in Table S1.

### Mice

Six-week-old female BALB/cOlaHsd mice were purchased from Envigo and housed in a BSL-3 facility under standard conditions (six mice per cage) with ad libitum access to food and water. FcγR/FcRn humanized mice were generated and characterized previously^51^, and maintained at the Comparative Bioscience Center at the Rockefeller University at a controlled ambient temperature environment with 12-h dark/light cycle.

### Cell lines

All mammalian cell lines were grown at 37°C and supplemented with a mixture of antibiotics (100 U/mL penicillin and 100 μg/mL streptomycin). Human embryonic kidney HEK293T cells were cultured in Dulbecco’s Modified Eagle Medium (DMEM) 10% FBS at 5% CO_2_. Expi293F cells were cultured in Expi medium (Gibco, A1435101) at 8% CO_2_. Human hepatocyte Huh-7.5 cells^82^ were cultured in DMEM supplemented with 10% FBS at 5% CO_2_. Human lung epithelial A549 cells (ATCC CRM-185) and A549-based AIR cells^48^ were cultured in DMEM 10% FBS at 5% CO_2_.

### Viruses

Eg-101, belonging to WNV lineage I, was originally isolated from human serum in Egypt in 1951,^83^ and was provided by the Collection of Arboviruses, Institute of Parasitology, Biology Centre of the Czech Academy of Sciences, Ceské Budějovice, Czech Republic. WNV strain 13-104, a representative of lineage II, was isolated from a *Culex modestus* mosquito in the Czech Republic in 2013^84^ and was kindly provided by Professors Zdeněk Hubálek and Ivo Rudolf, Institute of Vertebrate Biology, Czech Academy of Sciences, Valtice, Czech Republic. MVEV strain UVE/MVEV/UNK/AU/3329 (EVAg accession no. 001v-EVA145), SLEV strain UVE/SLEV/UNK/US/MSI-7 (EVAg accession no. 001v-EVA128), and JEV strain UVE/JEV/2009/LA/CNS769 were obtained from the EVAg and propagated in C6/36 prior to use.

### Blood sample collection and serum extraction

After informed consent was obtained, 3 mL of blood was collected in BD Vacutainer® SST™ Tubes (BD, Franklin Lakes, NJ, United States). Blood samples were allowed to clot at room temperature and, after centrifugation at 2000 × g for 10 min, serum was extracted and stored at − 80 °C until further use. Prior to use, sample were heat-inactivated (56°C for 1h) and then stored at 4°C thereafter. Orthoflavivirus naïve serum samples were obtained from healthy individuals with no history of orthoflavivirus infection or vaccination, and no IgG seroreactivity to WNV_EDIII_ protein in preliminary ELISA.

### Peripheral blood mononuclear cells (PBMCs) donor selection, large blood sampling and cell purification

Study participants were contacted during convalescence, starting from those with the highest serum neutralizing activity by the WNV_RVP_ assay, and invited to provide a large blood sample. Exclusion criteria for this part of the study were the same as those used for routine blood donors. Briefly, individuals with systemic diseases including diabetes and cardiovascular diseases, those aged below 18, or those with hemoglobin concentration <125 g/L (female) and <135 g/L (male) were excluded from the study. A detailed list of exclusion criteria (i.e., contraindications for blood donation) can be found in^85^. After determining hemoglobin levels (using the HemoCue Hb 301 System, HemoCue AB, Angelholm, Sweden), the candidates completed a blood donation questionnaire, were interviewed by a physician and underwent physical examination to establish suitability for large blood sampling. If successful up to this point of the selection, the candidate’s blood was additionally tested for (i) complete blood count (Celltac Alpha MEK-6500K automated hematology analyzer, Nihon Kohden, Tokyo, Japan); (ii) coagulation status (i.e., prothrombin time, international normalized ratio, and activated partial thromboplastin time; on a Siemens BCS XP fully automated hemostasis analyzer, Siemens Healthcare GmbH, Erlangen, Germany); (iii) levels of total proteins, albumins, immunoglobulins (isotypes G, M, and A), alanine aminotransferase, aspartate aminotransferase, gamma-glutamyl transferase, total and direct bilirubin, urea, creatinine, and C-reactive protein using spectrophotometry on a Cobas Integra 400 plus fully automated analyzer (Roche Diagnostics International AG, Rotkreuz, Switzerland). In case of satisfactory findings (i.e., results within reference values), the consenting convalescent study participant was enrolled as PBMC donor and 350mL of blood collected by venipunction (Double Blood Bag CPDA 350mL, Terumo Penpol Pvt Limited, India). The large blood sampling occurred approximately 10 months after hospital discharge (range: 9-11 months). Extraction of PBMCs was initiated within 5 hours after blood donation and was performed by gradient centrifugation using Histopaque. PBMCs were aliquoted in 90% Fetal Bovine Serum (FBS) and 10% dimethyl sulfoxide (DMSO) and stored in liquid nitrogen as previously described^86^, until further processing.

### Envelope domain III (EDIII) protein expression, purification and biotinylation

EDIII antigens were expressed in *E. coli* Rosetta 2 cells (DE3), purified from inclusion bodies, and refolded as previously detailed.^32^ Briefly, protein expression was induced with 1 mM isopropyl β-D-1-thiogalactopyranoside at 30°C for 4 hours. Cells were lysed, and the insoluble fraction containing inclusion bodies solubilized and refolded in a buffer containing 400 mM L-arginine, 100 mM Tris-base (pH 8.0), 2 mM ethylenediaminetetraacetic acid (EDTA), 0.2 mM phenylmethylsulfonyl fluoride, 5 mM reduced glutathione, 0.5 mM oxidized glutathione at 4°C. Refolded proteins were purified via size exclusion chromatography using a Superdex 75 column (Cytiva) in PBS buffer. The EDIII proteins were finally concentrated to 10–20 mg/ml. Codon-optimized EDIII expression plasmids corresponding to Orthoflaviviruses transmitted by *Culex* spp. mosquitoes (WNV lineage I, JEV, MVEV, SLEV and USUV), by ticks (POWV, KFDV, LGTV, LIV and OHFV) as well as by *Aedes* spp. mosquitoes (DENV serotypes 1-4, ZIKV, YFV) were previously reported.^32,34,79^ The plasmid to express the EDIII of WNV lineage II (residues 590– 690 of GenBank: UTQ11699.1) was obtained by site-directed mutagenesis of the plasmid encoding for the EDIII of WNV lineage I using the QuikChange Multi Site-Directed Mutagenesis Kit (Agilent Technologies) following manufacturer instructions. For the purification of antigen-specific memory B cells, the WNV lineage I EDIII was modified to include an Avi-tag following the C-terminal 6XHis-tag. The Avi-tag enabled site-directed biotinylation using the biotin-protein ligase standard reaction kit (BirA500), according to the manufacturer’s instructions (Avidity).

### Recombinant human IFN-α2 production and purification

An expression plasmid encoding human IFN-α2 was generated by cloning the coding sequence of human IFN-α2, fused at the C-terminus to an 8×His tag and AviTag™ into the pcDNA3.1 mammalian expression vector. Plasmid transfection was performed using linear polyethyleneimine hydrochloride (PEI-MAX; MW 40,000; Polysciences, 24765). For 300 mL Expi293F cell culture, 300 μg of plasmid DNA was mixed with 3 mg of PEI-MAX in 30 mL of OPTI-MEM medium (Gibco, 51985026) and incubated for 20 minutes at room temperature before being added dropwise to the cell suspension. On day 2 post-transfection, 100 mL of Expi293 expression medium (Gibco, A1435101) supplemented with 2 mL of soy hydrolysate (Sigma-Aldrich, 58903C) and 0.67 mL of D-Glucose (Sigma-Aldrich, G8769) was added. On day 4 post-transfection, cells were pelleted by centrifugation at 2500 x g for 30 min at 4°C, and the clarified supernatant was collected. It was then adjusted with 50 mM Tris-HCl (pH 7.5), 5 mM NiSO_4_, and 350 mM NaCl, filtered through 0.2 μm membranes, and loaded onto a 2 mL Ni-NTA affinity column (Thermo Fisher Scientific, 88222) pre-equilibrated with 10 mL equilibration buffer (50 mM Tris-HCl, pH 7.5, 350 mM NaCl). The resin was then washed with 50 mL wash buffer (50 mM Tris-HCl, pH 7.5, 500 mM NaCl, 40 mM Imidazole) and bound protein was eluted with elution buffer (50 mM Tris-HCl, pH 7.5, 200 mM NaCl, 250 mM Imidazole) in 10–15 fractions of 1 mL each. Fractions with the highest protein concentration, estimated by Bradford assay, were pooled and concentrated to 2 mL using a Vivaspin 20 centrifugal concentrator (10 kDa MWCO; Sigma-Aldrich, Z614653). The concentrated eluate was filtered (0.2 μm) and further purified by size-exclusion chromatography using a Superdex S200 16/60 column (Cytiva) pre-equilibrated with 1x PBS. After quality control analysis, the purified protein was aliquoted and stored at -80°C until use.

### ELISA

Detection of serum IgG or monoclonal antibody binding to EDIII antigen was measured by enzyme-linked immunosorbent assay (ELISA) using clear flat-bottom 384-well plates (Thermo Fisher Scientific, 464718), which were coated overnight at room temperature with 50 ng/well of EDIII protein in Phosphate-Buffered Saline (PBS). After washing with PBS-T (PBS + 0.05% Tween-20), plates were blocked with PBS, 1mM EDTA, 0.05% Tween-20 and 1% Bovine Serum Albumin (BSAlb) for 2 h at room temperature. After washing with PBS-T, serum or monoclonal antibodies (prediluted in PBS-T) were added to the plate and incubated for 1 h at room temperature. After washing with PBS-T, secondary sheep anti-human-IgG conjugated with horseradish Peroxidase (HRP; VWR, NA933; diluted 1:5000 in PBS-T) was added to the plate and incubated for an additional 1 h at room temperature. After the final washes, TMB (3,3⍰,5,5⍰-Tetramethylbenzidine) substrate (Thermo Fisher Scientific, 34021) was added to develop the plate, 1M sulfuric acid was added to stop the reaction, and the plates read at 450nm at the luminometer (BioTek, EPOCH2 microplate reader). Sera were serially diluted 3-fold, starting at 1:150 (Figure 1A). Recombinant monoclonal antibodies were serially diluted 3-fold, starting at 1.5 μg/mL (Figure 2A). Nonlinear regression analysis was used to determine the half-maximal effective concentration for binding (EC_50_) using Prism software (GraphPad), as previously detailed.^87^ For testing the crossreactivity to the EDIII of tick- or *Aedes* mosquito-borne Orthoflaviviruses, recombinant monoclonal antibodies were tested at 1.5 μg/mL according to the protocol described above. An anti-SARS CoV-2 monoclonal antibody, hr2.024 was used as isotype control.^86^ The broadly reactive EDIII antibody P003 was used as positive control.^79^

For the polyreactivity assay, 384-well plates were coated overnight at room temperature with 10 µg/ml for ssDNA, dsDNA, LPS, and KLH, and with 5 µg/ml for insulin in PBS. After washing with PBS +0.001% Tween-20-0.001% (PBSt), plates were blocked with PBS 1x, 0.5M EDTA, 0.0005% Tween-20 and 2.5% BSAlb for 2 h at room temperature. After washing with PBSt, monoclonal antibodies were added and incubated for 1 h. After washing with the same buffer, the same secondary antibody was added and incubated for 1 h. The development and stop of the reaction were performed as described above. ED38 polyreactive antibody^80^ was used as a positive control and ZIKV antibody Z021 was used as negative control.^32^

To determine the half-life of monoclonal antibody W010, human IgG levels were quantified in mouse serum samples by ELISA following previously published protocols. Briefly, high-binding, half-area 96-well microtiter plates (Nunc) were coated overnight at 4°C with neutravidin (3 μg/mL in PBS). After washing in PBS-T buffer, plates were incubated with biotinylated goat anti-human IgG antibodies for 1 h (5 μg/mL) and blocked for 2 h with PBS/2% bovine serum albumin. Serially diluted serum samples were incubated for 1 h, followed by incubation with horseradish peroxidase-conjugated anti-human IgG (1:10,000). Plates were developed using the TMB two-component peroxidase substrate kit (KPL), and reactions were stopped with the addition of 1 M sulfuric acid. Absorbance at 450 nm was immediately recorded using a SpectraMax Plus spectrophotometer (Molecular Devices), and background absorbance from negative control samples was subtracted. Antibody half-life was calculated as described.^88^

For the detection of serum autoantibodies against IFN-α2 (Figure S1P), an in-house developed ELISA was performed using the same protocol as for the EDIII, except that plates were coated with 5 μg/mL of human IFN-α2 recombinant protein and serum reactivity was tested starting at 1:50 dilution, followed by 4-fold serial dilutions.

### Determination of Tm

The melting temperature (Tm) of IgG antibodies (diluted in PBS) was determined using the ThermalShift Assay kit (ThermoFisher) on a QuantiStudio 6 flex thermal cycler, following manufacturer recommendations. Raw fluorescent values from quadruplicate samples were analyzed using the TSA craft tool as described^89^ and Tm (°C) are reported.

### Detection of anti-IFN-α2 and anti-IFN-ω autoantibodies

Detection of anti-IFN-α2 and anti-IFN-ω IgG (Figure 1D to F) was performed using a multiplexed bead-based assay essentially as described previously.^46,47^ Serum was tested at a 1:100 dilution. For each sample, median fluorescence intensity (MFI) values from the IFN-α2 or IFN-ω coated beads were obtained and made relative to the corresponding MFI value obtained from non-coated empty beads (fold over empty, FOE). Values were then normalized to in-house plate controls, and sera exhibiting normalized FOE values >5 SDs above the mean values obtained from a negative sample panel were considered positive for the specific anti-IFN-α2 or -ω IgG. Neutralizing anti-IFN-α2 or anti-IFN-ω activity was determined using the highly sensitive AIR cell system.^48^ Briefly, AIR cells were seeded at 30,000 cells per well in a 96-well plate in DMEM supplemented with 10% FBS, 100 U/mL penicillin and 100 µg/mL streptomycin (Gibco, 15140-122), and incubated overnight at 37°C, 5% CO_2_. Serum samples were diluted 1:20 in OptiMEM (Optimized Minimal Essential Medium; Gibco, 31985070) containing 0.01 ng/mL IFN-α2 (Novus Biologicals, NBP2-34971) or 0.02 ng/mL IFN-ω (Novus Biologicals, NBP2-35893), together with a live-cell Renilla luciferase substrate (EnduRen, Promega, E6481; 1:10,000), and incubated for 1 hour with constant shaking at 600 rpm. The serum:IFN-I mixtures were then used to stimulate AIR cells for 16 hours at 37°C, 5% CO_2_. Renilla luciferase activity levels were determined using a PerkinElmer EnVision plate reader (EV2104) and obtained luciferase values were normalized to in-house plate controls. Thresholds for positive neutralization activity were defined as a 75% reduction of IFN-I-induced luciferase value relative to the average value obtained from a negative sample panel.

### Single cell sorting

Isolation of WNV_EDIII_ memory B cells was performed as described.^34^ Briefly, the Pan B-cell isolation kit (Milteny Biotech, no. 130-101-638) and LS magnetic columns (Milteny Biotech; 130-042-401) were used to enrich B cells from PBMCs of WND convalescent individuals according to manufacturer’s instructions. The enriched B cells were subsequently incubated for 30 min on ice in FACS buffer (PBS, 2% FBS) with biotinylated WNV_EDIII_ conjugated to streptavidin-PE (eBioscience, Cat. No. 12-4317-87), with biotinylated WNV_EDIII_ conjugated to streptavidin-Alexa Fluor 647 (BioLegend; Cat. No. 405237) and with biotinylated

Ovalbumin (Sigma-Aldrich; A5503-1G) conjugated to streptavidin-BV711 (BD Horizon; 563262). The incubation was performed in the presence of anti-CD14-APC-eFluor 780 (Thermo Fischer Scientific, 47– 0149-42), anti-CD16-APC-eFluor 780 (Thermo Fischer Scientific, 47–0168-41), anti-CD8-APC-eFluor 780 (Invitrogen, 47–0086-41), anti-CD3-APC-eFluor 780 (Thermo Fischer Scientific, 47–0037-42), Zombie NIR™ (BioLegend, 423105) and anti-CD20-PE-Cy7 (BD Biosciences, 335828). Zombie^-^ NIR^−^CD14^−^CD16^−^CD3^−^CD8^−^CD20^+^Ova^−^WNV_EDIII_ -PE^+^ WNV_EDIII_ -AF647^+^ B cells were then single-cell sorted into 96-well plates containing lysis buffer (0.5× PBS, 10 mM dithiothreitol, and RNasin ribonuclease inhibitors; Promega, N2615) using the FACSymphony S6 cell sorter (BD Biosciences) and the analysis performed with FlowJo software. Sorted cells were snap-frozen on dry ice and then stored at −80°C.

### Antibody gene sequencing, cloning and expression

Single-cell RNA was reverse-transcribed (SuperScript III Reverse Transcriptase, Invitrogen 18080-044) and the resulting cDNA stored at -80°C until PCR-amplification of the variable Ig heavy and light genes by nested PCR reactions and Sanger sequencing, as detailed previously.^90,91^ Amplicons were next used for Sequence- and Ligation-Independent Cloning (SLIC) into antibody and Fab expression vectors.^32^ Recombinant monoclonal antibodies and Fabs were produced, purified and quality controlled as previously described.^32^ The E16 humanized sequence was retrieved from the literature^41^ and the antibody produced in-house starting from synthetic DNA (GenScript). Human one-armed IgG1 bispecific antibody was engineered using the variable regions of W010 or E16 and SARS-CoV-2 antibody sd1.040 in the CrossMAb format. Briefly, four expression plasmids were co-transfected at a 1:1:1:1 ratio, and antibodies were purified from cell supernatants as previously described.^86^

### Sequence analysis

Antibody sequences were analysed using in-house R and Perl scripts (available on GitHub;https://github.com/stratust/igpipeline). Briefly, sequences were annotated using Igblast tool v2; the sequences from the same cell were paired and assigned clonotypes based on V and J genes. Additionally, nucleotide somatic hypermutation and CDR3 length were analysed. Hypermutation analysis was based on the closest germlines sequences in IgBlast tool.

### Surface plasmon resonance (SPR) and epitope mapping

SPR assays were performed on a Biacore 8K instrument (Cytiva) at 25 °C, using 10 mM HEPES (4-(2-hydroxyethyl)-1-piperazineethanesulfonic acid) pH 7.4, 150 mM NaCl, 3 mM EDTA and 0.005% Tween-20 as running buffer. Kinetics parameters were determined as described.^92^ Antibodies were immobilized (150 nM) on the surface of CM5 chips (Cytiva) and increasing concentrations of the WNV_EDIII_ protein (15.62, 31.25, 62.5, 125, 250, 500 nM) were injected using a single-cycle kinetics setting; binding responses were corrected for unspecific binding and buffer responses. Curve fitting, data analysis and graphs were done with Biacore Insight Evaluation Software v.2.0.15.12933. Antibody competition experiments were performed to assess whether the antibodies recognize overlapping epitopes on the WNV_EDIII_ protein. Antibodies (W007, W009, W010, W014, W037, W048, W049) were immobilized as before and the WNV_EDIII_ protein was injected at a saturating concentration (500 nM). Immediately after, the second antibody was injected (150 nM) followed by a regeneration step with H_3_P0_4_ (0.1 M) to remove the EDIII protein and the second antibody. Additional cycles of injection of the EDIII protein, antibodies and H_3_P0_4_ were subsequently done. Curve fitting, data analysis and graphs were done with Biacore Insight Evaluation Software v.2.0.15.12933.

### Plasmids for the production of reporter virus particles (RVPs)

A WNV subgenomic replicon-expressing plasmid encoding Renilla luciferase (pWNVII-Rep-REN-IB; obtained from Ted Pierson at National Institutes of Health, NIH); ^93^ was modified by replacing Renilla by the NanoLuc luciferase coding sequence to generate pWNVII-Rep-NanoLuc. More in detail: plasmid pHIV-1_NL4-3_ΔEnv-NanoLuc^94^ was used as template for PCR with primers 5’-GTACGCGTATGGTCTTCACACTCGAAGA-3’ and 5’-GAAGTTACGCGCCGTCGCCAGAATGCGTTCGCACA-3’, while a second PCR was performed on plasmid pWNVII-Rep-REN-IB using primers 5’-GAACGCATTCTGGCGACGGCGCGTAACTTCGACCT-3’ and 5’-CACCCCACAGAGTGTGGGTT-3’. Upon purification, the two PCR products were combined by overlap PCR and ligated into plasmid pWNVII-Rep-REN-IB, previously digested with MluI (New England Biolabs, R3198S) and DraIII (New England Biolabs, R3510S), resulting in plasmid pWNVII-Rep-NanoLuc. The plasmid expressing the WNV CprME was previously reported (pWNV/TX02/CprME).^95^ For the generation of plasmids expressing the CprME of JEV (SA-14 strain; GenBank: M55506.1), MVEV (MK6684 strain; GenBank: KF751869.1), SLEV (Parton strain; GenBank: EF158070.1) and USUV (Vienna 2001 strain; GenBank: NC_006551), synthetic DNA segments with the corresponding CprME coding sequences were amplified by PCR prior to cloning into the NotI/PacI sites of pPOWV-LB-CprME^34^ resulting in plasmids pJEV-SA-14-CprME, pMVEV-MK6684-CprME, pSLEV-Parton-CprME and pUSUV-Vienna2001-CprME, respectively. The plasmids used to express the CprME of ZIKV and TBEV were previously reported.^32,34^

### RVP production

6-well plates were seeded with 7.5×10^5^ HEK293T cells/well. After 24 h cells were co-transfected using Lipofectamine 2000 (Invitrogen, 11668019) with a mix of pWNV-NanoLuc and CprME expressing plasmids at a 1:3 ratio.^34^ After 5 h of incubation at 37°C, the medium was replaced with DMEM high glucose GlutaMAX (Gibco) supplemented with 3% FBS, 100 U/mL penicillin and 100 µg/mL streptomycin. For the next 72 h, every 24 h RVPs were harvest, filtered (0.22 μm), aliquoted and frozen at -80°C, and the medium replaced. Infectivity was determined by titration (see Figure S1C).

### RVP neutralization assays

96-well plates were seeded with 1.5×10^4^ Huh-7.5 cells/well in 100 μL DMEM high glucose GlutaMAX (Gibco) supplemented with 10% FBS, 1% sodium pyruvate, 1% non-essential amino acids, 100 U/mL penicillin and 100 µg/mL streptomycin. After 24 h, the antibody containing samples were diluted with Ba-1 medium (Medium 199 (Gibco), 1% BSAlb, 100 U/mL penicillin and 100 µg/mL streptomycin), then 200 μL diluted serum or antibody were combined with 200 μL diluted RVPs diluted in Ba-1 medium and incubated for 1 h at 37°C. 100 μL of the mix were then added in triplicate to the plate and incubated overnight at 37°C. The supernatant was next aspirated, the cells washed with PBS, and 50 μL lysis buffer added (Promega, E1531). The Nano-Glo Luciferase Assay System (Promega, N1120) with GloMAx Discover System reader (Promega) were used to measure the NanoLuc luciferase activity of the lysate. Serum was diluted 1:100 for the WNV_RVP_ neutralization screening (Figure 1B). Otherwise, it was diluted 1:150 and then serially 1:3. Recombinant monoclonal antibodies were tested starting with 10 μg/mL (WNV RVPs) or 1 μg/mL (JEV, MVEV, SLEV and USUV RVPs), Fab molecules at 6.7 µg/mL, and then serially diluted 1:3. Nonlinear regression analysis was used to determine the NT_50_ and IC_50_ values using Prism software (GraphPad). In the cross-neutralization screening against the panel of orthoflavivirus RVPs (Figure S6A), recombinant antibodies were tested at a single concentration of 500 ng/mL using the protocol described above, and the results compared to no antibody control.

### Plaque assay

The viral titers were determined using a plaque assay. Viruses were serially diluted in DMEM and incubated with A549 cells (2 × 10^5^ per well) for 3 hours at 37⍰°C in 5% CO_2_. Cells were then overlaid with DMEM containing 1.5% carboxymethylcellulose and incubated under the same conditions for either 4 days (WNV lineage I and II) or 5 days (JEV, MVEV and SLEV) prior to washing with PBS, and staining with naphthalene black solution to visualize plaques. The viral titers were calculated and expressed as plaque-forming units per milliliter (PFU/mL).

### Virus neutralization assay and immunofluorescent staining

All tested monoclonal antibodies were initially diluted in DMEM medium to a starting concentration of 10⍰µg/mL and subsequently subjected to a 2-fold serial dilution in 96-well plates. Following dilution, 50 PFU of WNV, lineage I or II, were added to each well and the mixtures were incubated for 90 minutes at 37⍰°C in 5% CO_2_. After incubation, 3 × 10^4^ A549 cells were added to each well, and the plates were further incubated under the same conditions. Cytopathic effects were monitored five days post-infection using an inverted microscope. Virus neutralization assay was performed in octuplicates for each concentration and the result expressed as the percentage of neutralization over no-antibody control for each concentration (Figure 3B).

A similar procedure was employed for the fluorescence-based neutralization assay (Figure S3A). Three (JEV, MVEV and SLEV) or four days (WNV lineage I and II) post-infection, cells were fixed with cold acetone/methanol (1:1) and subsequently blocked with 10% FBS. Cells were then incubated with a mouse anti-flavivirus antibody in 1:250 dilution (Sigma, MAB10216), washed three times with PBS-T, and incubated with a fluorescently labelled secondary goat anti-mouse antibody in 1:500 dilution (Invitrogen, A32723). After three additional washes with PBS-T, cell nuclei were stained using DAPI at a 1:2000 dilution (Sigma, D9542). Fluorescence images were acquired using the ImageXpress Pico automated imaging system and CellReporterXpress software (Molecular Devices, USA).

### Selection of antibody-resistant WNV variants

To select for antibody-resistant variants of W NV 13-104, the virus was serially passaged in the presence of increasing concentrations of the monoclonal antibody W010 in A549 cells. Briefly, W010 was diluted in DMEM and incubated with 5000 PFU of WNV 13-104 for 90 minutes at 37⍰°C in a 24-well plate. Following incubation, 2 × 10^5^ A549 cells were added to each well. In parallel, wild-type virus was passaged under identical conditions without the addition of antibody as a control. Five days post-infection, supernatants from wells displaying CPE were harvested and used for subsequent passages with progressively higher concentrations of W010. After RNA isolation (QIAGEN, 52904), viral genomic RNA was quantified at each passage using quantitative reverse transcription PCR with the Advanced Kit for West Nile Virus (Genesig, Z-Path-WNV-EASY) and Lyophilised OneStep qRT-PCR Master Mix (Oasig, Z-oasig-onestep-150), according to the manufacturers’ instructions. The virus envelope (E) gene was sequenced with primers WNV_1A (CCTGGTGGCACCAGCATACAGC) + WNV_1B (GTGGTGCTTCCAGCACTGCTCC) and WNV_2A (GCGAAGTCCTTCCTGGTTCACCGA) + WNV_2B (CACAGCCTGTGTCAGCATGGACG) and analyzed from viral populations passaged in the presence of 50⍰ng/mL W010 to assess mutations associated with antibody resistance.

The plasmid to express the mutant WNVRVP N368T was obtained by site-directed mutagenesis of the plasmid pWNV/TX02/CprME using the QuikChange Multi Site-Directed Mutagenesis following manufacturer instructions.

### Structure determination via X-ray crystallography

W037 Fab and W014 Fab were expressed in Expi293F cells, and subsequently purified via Nickel-NTA (nitriloacetic acid) as previously described.^34^ W010 IgG was expressed in Expi293F cells, purified by Protein A (ProA) affinity chromatography and cleaved with 1.5% papain to generate Fab fragments. Fabs were subsequently isolated from Fc components or uncleaved IgG by purification over a ProA column. All Fabs were purified via size exclusion chromatography (SEC) over a Superdex 200 10/300 column against 1x TBS (20 mM Tris pH 8.0, 150 mM NaCl). Fab-WNV EDIII complexes were generated by combining WNV EDIII with Fab at a 2:1 ratio overnight at room temperature, followed by purification via SEC using a Superdex 200 10/300 column to separate complex from excess WNV EDIII. Fractions attributed to the complex were analyzed by Sodium Dodecyl Sulfate-Polyacrylamide Gel Electrophoresis (SDS-PAGE) prior to pooling and concentrating to 15 mg/mL.

Crystallization trials were set up using the sitting drop vapor diffusion method by mixing equal volumes of complex and reservoir using a Mosquito LCP liquid handling robot (SPT LabTech) and commercially available 96-well crystallization screens (Hampton Research). Crystals were grown at 22 °C and observed in multiple conditions. The single crystal used for structure determination of W037 Fab-WNV EDIII was obtained in 0.2 M magnesium chloride hexahydrate, 20% w/v polyethylene glycol (PEG) 3,350. The single crystal used for structure determination of W010 Fab-WNV EDIII was obtained in 20% w/v PEG4,000, 0.2M disodium malonate. The single crystal used for structure determination of W014 Fab-WNV EDIII was obtained in 20% w/v 2-propanol, 20% w/v PEGME (polyethylene glycol monomethyl ether) 2000, 0.1 MES (2-(n-morpholino) ethanesulfonic acid). The single crystal used for structure determination of W049 Fab-WNV EDIII was obtained in 12% w/v PEG3,350 and 0.1M disodium DL-malate. All crystals were cryoprotected in a solution matching the reservoir and 20% glycerol and then cryocooled in liquid nitrogen.

All X-ray diffraction data was collected at the Stanford Synchrotron Radiation Lightsource on beamline 12-2 with a Pilatus 6M detector at a wavelength of 0.979Å and temperature of 100K. Data from a single crystal was indexed and integrated in XDS^96^ and merged using AIMLESS in CCP4.^97^ Structures were determined using molecular replacement in Phenix PHASER^98^ using each of the following individual chains as search models with CDR3 loops trimmed: WNV EDIII for all structures (PDB 6UTE); W037 Fab V_H_ (PDB 6Z3K) and V_L_ (PDB 6WOZ); W010 Fab V_H_ (PDB 6DWC) and V_L,_ (7BEI); W014 Fab V_H_ (PDB 5GGU) and V_L_ (PDB 7BEI); W049 Fab V_H_ and V_L_ (PDB 6DWC). Coordinates were refined using iterative rounds of automated and interactive refinement in Phenix^99^ and Coot,^100^ respectively. Statistics for the final models can be found in table S3.

### Structural Analyses

Complementarity-determining regions (CDRs) and somatic mutations were annotated using IMGT (International ImMunoGeneTics information system) V-QUEST.^101^ Structural visualizations were generated with ChimeraX.^102^ Buried surface area calculations were performed via the PDBePISA web server.^103^ Residue contacts were defined as atom-atom distances under 4 Å between different chains. Hydrogen bonds were identified using a distance cutoff of 3.5 Å and an angle criterion with an A–D–H angle exceeding 90°. Root-mean-square deviation (RMSD) values were calculated using ChimeraX. Antibody residue numbering follows the Kabat scheme. The atomic models and corresponding structure factor files for all structures are available in the Protein Data Bank (PDB) under the following accession codes: W010 Fab-WNV EDIII (PDB 9ZRM), W014 Fab-WNV EDIII (PDB 9ZRN), W037 Fab-WNV EDIII (PDB 9ZRO), W049 Fab-WNV EDIII (PDB 9ZRP).

### Mouse protection experiments

The animal experiments adhered to all relevant European Union guidelines for animal research and were conducted in accordance with Czech national law (Animal Welfare Act No. 246/1992 Coll.), ensuring the ethical use and protection of experimental animals. The study protocol was approved by the Ministry of Agriculture of the Czech Republic (permit nos. 22006/2016-MZe-17214 and MZE-23932/2024-13143).

Five experimental groups (n=12 per group) were established. Mice were inoculated intraperitoneally with 200 μL PBS containing monoclonal antibody W010 (30 µg/mouse) either one day before infection or 1, 3, or 5 days post-infection. Isotype control was antibody 10-1074.^104^ On the day of infection, all mice were challenged subcutaneously with a lethal dose of WNV strain 13-104 (10^4^ PFU/mouse). Mouse protection experiments with W014 were conducted as with W010, except that mice were treated only at day -1 and with 300 µg of either W014 (n=10) or isotype control 10-1074 (n=5).

For mouse protection experiments in the presence of anti-IFNAR antibodies, four experimental groups (n=6 per group) were established. Three groups were inoculated intraperitoneally with 200 μl PBS containing 100 μg anti-IFNAR1 antibody (BioXCell, #BP0241), while the control group received 200⍰μL of PBS alone. The anti-IFNAR antibody or PBS alone were administered one day before infection and then every other day, for a total of three doses. On the day of infection, the control group and one of the anti-IFNAR-treated groups were intraperitoneally injected with 200⍰μL of PBS containing an isotype control monoclonal antibody. The other 2 anti-IFNAR-treated groups received 200⍰μL of PBS containing 200⍰μg of W010 on either day 0 or 1 after infection. All mice were challenged subcutaneously with a lethal dose (10^4^ PFU per mouse) of WNV strain 13-104.

The animals were closely monitored for health status, clinical symptoms, and survival for up to 18 or 28 days. Survival rates were analyzed by log rank Mantel-Cox test in GraphPad Prism.

## QUANTIFICATION AND STATISTICAL ANALYSIS

Statistical tests were as indicated throughout the methods and in the figures or their legends. The analyses were performed with Prism software (version 10.6.1; GraphPad), unless otherwise noted.

## REFERENCES

1. Gould, E.A., and Solomon, T. (2008). Pathogenic flaviviruses. Lancet 371, 500–509. 10.1016/S0140-6736(08)60238-X.

2. Dutta, S.K., and Langenburg, T. (2023). A Perspective on Current Flavivirus Vaccine Development: A Brief Review. Viruses 15. 10.3390/v15040860.

3. Pierson, T.C., and Diamond, M.S. (2020). The continued threat of emerging flaviviruses. Nat Microbiol 5, 796–812. 10.1038/s41564-020-0714-0.

4. Martin-Acebes, M.A., and Saiz, J.C. (2012). West Nile virus: A re-emerging pathogen revisited. World J Virol 1, 51–70. 10.5501/wjv.v1.i2.51.

5. Kramer, L.D., Li, J., and Shi, P.Y. (2007). West Nile virus. Lancet Neurol 6, 171–181. 10.1016/S1474-4422(07)70030-3.

6. Murray, K., Baraniuk, S., Resnick, M., Arafat, R., Kilborn, C., Cain, K., Shallenberger, R., York, T.L., Martinez, D., Hellums, J.S., et al. (2006). Risk factors for encephalitis and death from West Nile virus infection. Epidemiol Infect 134, 1325–1332. 10.1017/S0950268806006339.

7. Iwamoto, M., Jernigan, D.B., Guasch, A., Trepka, M.J., Blackmore, C.G., Hellinger, W.C., Pham, S.M., Zaki, S., Lanciotti, R.S., Lance-Parker, S.E., et al. (2003). Transmission of West Nile virus from an organ donor to four transplant recipients. N Engl J Med 348, 2196–2203. 10.1056/NEJMoa022987.

8. Gervais, A., Rovida, F., Avanzini, M.A., Croce, S., Marchal, A., Lin, S.C., Ferrari, A., Thorball, C.W., Constant, O., Le Voyer, T., et al. (2023). Autoantibodies neutralizing type I IFNs underlie West Nile virus encephalitis in approximately 40% of patients. J Exp Med 220. 10.1084/jem.20230661.

9. Lim, J.K., Glass, W.G., McDermott, D.H., and Murphy, P.M. (2006). CCR5: no longer a “good for nothing” gene--chemokine control of West Nile virus infection. Trends Immunol 27, 308–312. 10.1016/j.it.2006.05.007.

10. Lim, J.K., Louie, C.Y., Glaser, C., Jean, C., Johnson, B., Johnson, H., McDermott, D.H., and Murphy, P.M. (2008). Genetic deficiency of chemokine receptor CCR5 is a strong risk factor for symptomatic West Nile virus infection: a meta-analysis of 4 cohorts in the US epidemic. J Infect Dis 197, 262–265. 10.1086/524691.

11. Montgomery, R.R. (2017). Age-related alterations in immune responses to West Nile virus infection. Clin Exp Immunol 187, 26–34. 10.1111/cei.12863.

12. Gervais, A., Trespidi, F., Ferrari, A., Rovida, F., Marchal, A., Croce, S., Cassaniti, I., Moratti, M., Uhrlaub, J.L., Florian, D.M., et al. (2025). Autoantibodies neutralizing type I IFNs in 40% of patients with WNV encephalitis in seven new cohorts. medRxiv. 10.1101/2025.08.31.25334556.

13. Bastard, P., Michailidis, E., Hoffmann, H.H., Chbihi, M., Le Voyer, T., Rosain, J., Philippot, Q., Seeleuthner, Y., Gervais, A., Materna, M., et al. (2021). Auto-antibodies to type I IFNs can underlie adverse reactions to yellow fever live attenuated vaccine. J Exp Med 218. 10.1084/jem.20202486.

14. Gervais, A., Marchal, A., Fortova, A., Berankova, M., Krbkova, L., Pychova, M., Salat, J., Zhao, S., Kerrouche, N., Le Voyer, T., et al. (2024). Autoantibodies neutralizing type I IFNs underlie severe tick-borne encephalitis in approximately 10% of patients. J Exp Med 221. 10.1084/jem.20240637.

15. Gervais, A., Bastard, P., Bizien, L., Delifer, C., Tiberghien, P., Rodrigo, C., Trespidi, F., Angelini, M., Rossini, G., Lazzarotto, T., et al. (2024). Auto-Abs neutralizing type I IFNs in patients with severe Powassan, Usutu, or Ross River virus disease. J Exp Med 221. 10.1084/jem.20240942.

16. Lin, S.C., Zhao, F.R., Janova, H., Gervais, A., Rucknagel, S., Murray, K.O., Casanova, J.L., and Diamond, M.S. (2023). Blockade of interferon signaling decreases gut barrier integrity and promotes severe West Nile virus disease. Nat Commun 14, 5973. 10.1038/s41467-023-41600-3.

17. Riccardo, F., Bella, A., Monaco, F., Ferraro, F., Petrone, D., Mateo-Urdiales, A., Andrianou, X.D., Del Manso, M., Venturi, G., Fortuna, C., et al. (2022). Rapid increase in neuroinvasive West Nile virus infections in humans, Italy, July 2022. Euro Surveill 27. 10.2807/1560-7917.ES.2022.27.36.2200653.

18. Ciota, A.T., and Kramer, L.D. (2013). Vector-virus interactions and transmission dynamics of West Nile virus. Viruses 5, 3021–3047. 10.3390/v5123021.

19. Koch, R.T., Erazo, D., Folly, A.J., Johnson, N., Dellicour, S., Grubaugh, N.D., and Vogels, C.B.F. (2024). Genomic epidemiology of West Nile virus in Europe. One Health 18, 100664. 10.1016/j.onehlt.2023.100664.

20. Hadfield, J., Brito, A.F., Swetnam, D.M., Vogels, C.B.F., Tokarz, R.E., Andersen, K.G., Smith, R.C., Bedford, T., and Grubaugh, N.D. (2019). Twenty years of West Nile virus spread and evolution in the Americas visualized by Nextstrain. PLoS Pathog 15, e1008042. 10.1371/journal.ppat.1008042.

21. Farooq, Z., Sjodin, H., Semenza, J.C., Tozan, Y., Sewe, M.O., Wallin, J., and Rocklov, J. (2023). European projections of West Nile virus transmission under climate change scenarios. One Health 16, 100509. 10.1016/j.onehlt.2023.100509.

22. Danforth, M.E., Fischer, M., Snyder, R.E., Lindsey, N.P., Martin, S.W., and Kramer, V.L. (2021). Characterizing Areas with Increased Burden of West Nile Virus Disease in California, 2009-2018. Vector Borne Zoonotic Dis 21, 620–627. 10.1089/vbz.2021.0014.

23. Jabr, R., Khatri, A., Anderson, A.D., Garcia, L.C., Viotti, J.B., Natori, Y., Raja, M., Camargo, J.F., and Morris, M.I. (2023). Early administration of SARS-CoV-2 monoclonal antibody reduces the risk of mortality in hematologic malignancy and hematopoietic cell transplant patients with COVID-19. Transpl Infect Dis 25, e14006. 10.1111/tid.14006.

24. Sneller, M.C., Blazkova, J., Justement, J.S., Shi, V., Kennedy, B.D., Gittens, K., Tolstenko, J., McCormack, G., Whitehead, E.J., Schneck, R.F., et al. (2022). Combination anti-HIV antibodies provide sustained virological suppression. Nature 606, 375–381. 10.1038/s41586-022-04797-9.

25. Saxena, D., Kaul, G., Dasgupta, A., and Chopra, S. (2021). Atoltivimab/maftivimab/odesivimab (Inmazeb) combination to treat infection caused by Zaire ebolavirus. Drugs Today (Barc) 57, 483–490. 10.1358/dot.2021.57.8.3280599.

26. Low, J.G., Ng, J.H.J., Ong, E.Z., Kalimuddin, S., Wijaya, L., Chan, Y.F.Z., Ng, D.H.L., Tan, H.C., Baglody, A., Chionh, Y.H., et al. (2020). Phase 1 Trial of a Therapeutic Anti-Yellow Fever Virus Human Antibody. N Engl J Med 383, 452–459. 10.1056/NEJMoa2000226.

27. Gunale, B., Farinola, N., Kamat, C.D., Poonawalla, C.S., Pisal, S.S., Dhere, R.M., Miller, C., and Kulkarni, P.S. (2024). An observer-blind, randomised, placebo-controlled, phase 1, single ascending dose study of dengue monoclonal antibody in healthy adults in Australia. Lancet Infect Dis 24, 639–649. 10.1016/S1473-3099(24)00030-6.

28. Ltd., T.P. (2018). Safety and Tolerability of an Antibody Against Zika Virus (Tyzivumab) in Humans. https://clinicaltrials.gov/study/NCT03443830.

29. Beigel, J.H., Nordstrom, J.L., Pillemer, S.R., Roncal, C., Goldwater, D.R., Li, H., Holland, P.C., Johnson, S., Stein, K., and Koenig, S. (2010). Safety and pharmacokinetics of single intravenous dose of MGAWN1, a novel monoclonal antibody to West Nile virus. Antimicrob Agents Chemother 54, 2431–2436. 10.1128/AAC.01178-09.

30. Gnann, J.W., Jr., Agrawal, A., Hart, J., Buitrago, M., Carson, P., Hanfelt-Goade, D., Tyler, K., Spotkov, J., Freifeld, A., Moore, T., et al. (2019). Lack of Efficacy of High-Titered Immunoglobulin in Patients with West Nile Virus Central Nervous System Disease. Emerg Infect Dis 25, 2064–2073. 10.3201/eid2511.190537.

31. Kanai, R., Kar, K., Anthony, K., Gould, L.H., Ledizet, M., Fikrig, E., Marasco, W.A., Koski, R.A., and Modis, Y. (2006). Crystal structure of west nile virus envelope glycoprotein reveals viral surface epitopes. J Virol 80, 11000–11008. 10.1128/JVI.01735-06.

32. Robbiani, D.F., Bozzacco, L., Keeffe, J.R., Khouri, R., Olsen, P.C., Gazumyan, A., Schaefer-Babajew, D., Avila-Rios, S., Nogueira, L., Patel, R., et al. (2017). Recurrent Potent Human Neutralizing Antibodies to Zika Virus in Brazil and Mexico. Cell 169, 597–609 e511. 10.1016/j.cell.2017.04.024.

33. Dai, L., Song, J., Lu, X., Deng, Y.Q., Musyoki, A.M., Cheng, H., Zhang, Y., Yuan, Y., Song, H., Haywood, J., et al. (2016). Structures of the Zika Virus Envelope Protein and Its Complex with a Flavivirus Broadly Protective Antibody. Cell Host Microbe 19, 696–704. 10.1016/j.chom.2016.04.013.

34. Agudelo, M., Palus, M., Keeffe, J.R., Bianchini, F., Svoboda, P., Salat, J., Peace, A., Gazumyan, A., Cipolla, M., Kapoor, T., et al. (2021). Broad and potent neutralizing human antibodies to tick-borne flaviviruses protect mice from disease. J Exp Med 218. 10.1084/jem.20210236.

35. Dai, L., Wang, Q., Qi, J., Shi, Y., Yan, J., and Gao, G.F. (2016). Molecular basis of antibody-mediated neutralization and protection against flavivirus. IUBMB Life 68, 783–791. 10.1002/iub.1556.

36. Wang, Q., Yan, J., and Gao, G.F. (2017). Monoclonal Antibodies against Zika Virus: Therapeutics and Their Implications for Vaccine Design. J Virol 91. 10.1128/JVI.01049-17.

37. Malonis, R.J., Georgiev, G.I., Haslwanter, D., VanBlargan, L.A., Fallon, G., Vergnolle, O., Cahill, S.M., Harris, R., Cowburn, D., Chandran, K., et al. (2022). A Powassan virus domain III nanoparticle immunogen elicits neutralizing and protective antibodies in mice. PLoS Pathog 18, e1010573. 10.1371/journal.ppat.1010573.

38. Deng, W.L., Guan, C.Y., Liu, K., Zhang, X.M., Feng, X.L., Zhou, B., Su, X.D., and Chen, P.Y. (2014). Fine mapping of a linear epitope on EDIII of Japanese encephalitis virus using a novel neutralizing monoclonal antibody. Virus Res 179, 133–139. 10.1016/j.virusres.2013.10.022.

39. Stettler, K., Beltramello, M., Espinosa, D.A., Graham, V., Cassotta, A., Bianchi, S., Vanzetta, F., Minola, A., Jaconi, S., Mele, F., et al. (2016). Specificity, cross-reactivity, and function of antibodies elicited by Zika virus infection. Science 353, 823–826. 10.1126/science.aaf8505.

40. Throsby, M., Geuijen, C., Goudsmit, J., Bakker, A.Q., Korimbocus, J., Kramer, R.A., Clijsters-van der Horst, M., de Jong, M., Jongeneelen, M., Thijsse, S., et al. (2006). Isolation and characterization of human monoclonal antibodies from individuals infected with West Nile Virus. J Virol 80, 6982–6992. 10.1128/JVI.00551-06.

41. Oliphant, T., Engle, M., Nybakken, G.E., Doane, C., Johnson, S., Huang, L., Gorlatov, S., Mehlhop, E., Marri, A., Chung, K.M., et al. (2005). Development of a humanized monoclonal antibody with therapeutic potential against West Nile virus. Nat Med 11, 522–530. 10.1038/nm1240.

42. Knox, J., Cowan, R.U., Doyle, J.S., Ligtermoet, M.K., Archer, J.S., Burrow, J.N., Tong, S.Y., Currie, B.J., Mackenzie, J.S., Smith, D.W., et al. (2012). Murray Valley encephalitis: a review of clinical features, diagnosis and treatment. Med J Aust 196, 322–326. 10.5694/mja11.11026.

43. Ortiz-Martinez, Y., Vega-Useche, L., Villamil-Gomez, W.E., and Rodriguez-Morales, A.J. (2017). Saint Louis Encephalitis Virus, another re-emerging arbovirus: a literature review of worldwide research. Infez Med 25, 77–79.

44. Carson, P.J., Prince, H.E., Biggerstaff, B.J., Lanciotti, R., Tobler, L.H., and Busch, M. (2014). Characteristics of antibody responses in West Nile virus-seropositive blood donors. J Clin Microbiol 52, 57–60. 10.1128/JCM.01932-13.

45. Robbiani, D.F., Olsen, P.C., Costa, F., Wang, Q., Oliveira, T.Y., Nery, N., Jr., Aromolaran, A., do Rosario, M.S., Sacramento, G.A., Cruz, J.S., et al. (2019). Risk of Zika microcephaly correlates with features of maternal antibodies. J Exp Med 216, 2302–2315. 10.1084/jem.20191061.

46. Fernbach, S., Mair, N.K., Abela, I.A., Groen, K., Kuratli, R., Lork, M., Thorball, C.W., Bernasconi, E., Filippidis, P., Leuzinger, K., et al. (2024). Loss of tolerance precedes triggering and lifelong persistence of pathogenic type I interferon autoantibodies. J Exp Med 221. 10.1084/jem.20240365.

47. Busnadiego, I., Abela, I.A., Frey, P.M., Hofmaenner, D.A., Scheier, T.C., Schuepbach, R.A., Buehler, P.K., Brugger, S.D., and Hale, B.G. (2022). Critically ill COVID-19 patients with neutralizing autoantibodies against type I interferons have increased risk of herpesvirus disease. PLoS Biol 20, e3001709. 10.1371/journal.pbio.3001709.

48. Groen, K., Kuratli, R., Sar, L., Vasou, A., Huber, M., Hughes, D.J., and Hale, B.G. (2024). Highly sensitive reporter cell line for detection of interferon types I-III and their neutralization by antibodies. Eur J Immunol 54, e2451325. 10.1002/eji.202451325.

49. Bastard, P., Gervais, A., Le Voyer, T., Rosain, J., Philippot, Q., Manry, J., Michailidis, E., Hoffmann, H.H., Eto, S., Garcia-Prat, M., et al. (2021). Autoantibodies neutralizing type I IFNs are present in ∼4% of uninfected individuals over 70 years old and account for ∼20% of COVID-19 deaths. Sci Immunol 6. 10.1126/sciimmunol.abl4340.

50. Priyamvada, L., Cho, A., Onlamoon, N., Zheng, N.Y., Huang, M., Kovalenkov, Y., Chokephaibulkit, K., Angkasekwinai, N., Pattanapanyasat, K., Ahmed, R., et al. (2016). B Cell Responses during Secondary Dengue Virus Infection Are Dominated by Highly Cross-Reactive, Memory-Derived Plasmablasts. J Virol 90, 5574–5585. 10.1128/JVI.03203-15.

51. Borghi, S., Bournazos, S., Thulin, N.K., Li, C., Gajewski, A., Sherwood, R.W., Zhang, S., Harris, E., Jagannathan, P., Wang, L.X., et al. (2020). FcRn, but not FcgammaRs, drives maternal-fetal transplacental transport of human IgG antibodies. Proc Natl Acad Sci U S A 117, 12943–12951. 10.1073/pnas.2004325117.

52. Diamond, M.S., Pierson, T.C., and Fremont, D.H. (2008). The structural immunology of antibody protection against West Nile virus. Immunol Rev 225, 212–225. 10.1111/j.1600-065X.2008.00676.x.

53. Hardy, J.M., Newton, N.D., Modhiran, N., Scott, C.A.P., Venugopal, H., Vet, L.J., Young, P.R., Hall, R.A., Hobson-Peters, J., Coulibaly, F., and Watterson, D. (2021). A unified route for flavivirus structures uncovers essential pocket factors conserved across pathogenic viruses. Nat Commun 12, 3266. 10.1038/s41467-021-22773-1.

54. Reisen, W.K., Fang, Y., and Martinez, V.M. (2006). Effects of temperature on the transmission of west nile virus by Culex tarsalis (Diptera: Culicidae). J Med Entomol 43, 309–317. 10.1603/0022-2585(2006)043[0309:EOTOTT]2.0.CO;2.

55. Erazo, D., Grant, L., Ghisbain, G., Marini, G., Colon-Gonzalez, F.J., Wint, W., Rizzoli, A., Van Bortel, W., Vogels, C.B.F., Grubaugh, N.D., et al. (2024). Contribution of climate change to the spatial expansion of West Nile virus in Europe. Nat Commun 15, 1196. 10.1038/s41467-024-45290-3.

56. Klingelhofer, D., Braun, M., Kramer, I.M., Reuss, F., Muller, R., Groneberg, D.A., and Bruggmann, D. (2023). A virus becomes a global concern: research activities on West-Nile virus. Emerg Microbes Infect 12, 2256424. 10.1080/22221751.2023.2256424.

57. Isaacs, A., and Lindenmann, J. (1957). Virus interference. I. The interferon. Proc R Soc Lond B Biol Sci 147, 258–267. 10.1098/rspb.1957.0048.

58. Isaacs, A., Lindenmann, J., and Valentine, R.C. (1957). Virus interference. II. Some properties of interferon. Proc R Soc Lond B Biol Sci 147, 268–273. 10.1098/rspb.1957.0049.

59. Isaacs, A., and Burke, D.C. (1959). Viral interference and interferon. Br Med Bull 15, 185–188. 10.1093/oxfordjournals.bmb.a069760.

60. Pinto, A.K., Daffis, S., Brien, J.D., Gainey, M.D., Yokoyama, W.M., Sheehan, K.C., Murphy, K.M., Schreiber, R.D., and Diamond, M.S. (2011). A temporal role of type I interferon signaling in CD8+ T cell maturation during acute West Nile virus infection. PLoS Pathog 7, e1002407. 10.1371/journal.ppat.1002407.

61. Samuel, M.A., and Diamond, M.S. (2005). Alpha/beta interferon protects against lethal West Nile virus infection by restricting cellular tropism and enhancing neuronal survival. J Virol 79, 13350–13361. 10.1128/JVI.79.21.13350-13361.2005.

62. Daffis, S., Samuel, M.A., Suthar, M.S., Keller, B.C., Gale, M., Jr., and Diamond, M.S. (2008). Interferon regulatory factor IRF-7 induces the antiviral alpha interferon response and protects against lethal West Nile virus infection. J Virol 82, 8465–8475. 10.1128/JVI.00918-08.

63. Mathian, A., Breillat, P., Dorgham, K., Bastard, P., Charre, C., Lhote, R., Quentric, P., Moyon, Q., Mariaggi, A.A., Mouries-Martin, S., et al. (2022). Lower disease activity but higher risk of severe COVID-19 and herpes zoster in patients with systemic lupus erythematosus with pre-existing autoantibodies neutralising IFN-alpha. Ann Rheum Dis 81, 1695–1703. 10.1136/ard-2022-222549.

64. Sokal, A., Bastard, P., Chappert, P., Barba-Spaeth, G., Fourati, S., Vanderberghe, A., Lagouge-Roussey, P., Meyts, I., Gervais, A., Bouvier-Alias, M., et al. (2023). Human type I IFN deficiency does not impair B cell response to SARS-CoV-2 mRNA vaccination. J Exp Med 220. 10.1084/jem.20220258.

65. Bastard, P., Vazquez, S.E., Liu, J., Laurie, M.T., Wang, C.Y., Gervais, A., Le Voyer, T., Bizien, L., Zamecnik, C., Philippot, Q., et al. (2023). Vaccine breakthrough hypoxemic COVID-19 pneumonia in patients with auto-Abs neutralizing type I IFNs. Sci Immunol 8, eabp8966. 10.1126/sciimmunol.abp8966.

66. Keeffe, J.R., Van Rompay, K.K.A., Olsen, P.C., Wang, Q., Gazumyan, A., Azzopardi, S.A., Schaefer-Babajew, D., Lee, Y.E., Stuart, J.B., Singapuri, A., et al. (2018). A Combination of Two Human Monoclonal Antibodies Prevents Zika Virus Escape Mutations in Non-human Primates. Cell Rep 25, 1385–1394 e1387. 10.1016/j.celrep.2018.10.031.

67. Kaufmann, B., Nybakken, G.E., Chipman, P.R., Zhang, W., Diamond, M.S., Fremont, D.H., Kuhn, R.J., and Rossmann, M.G. (2006). West Nile virus in complex with the Fab fragment of a neutralizing monoclonal antibody. Proc Natl Acad Sci U S A 103, 12400–12404. 10.1073/pnas.0603488103.

68. Dowd, K.A., and Pierson, T.C. (2011). Antibody-mediated neutralization of flaviviruses: a reductionist view. Virology 411, 306–315. 10.1016/j.virol.2010.12.020.

69. Georgiev, G.I., Malonis, R.J., Wirchnianski, A.S., Wessel, A.W., Jung, H.S., Cahill, S.M., Nyakatura, E.K., Vergnolle, O., Dowd, K.A., Cowburn, D., et al. (2022). Resurfaced ZIKV EDIII nanoparticle immunogens elicit neutralizing and protective responses in vivo. Cell Chem Biol 29, 811–823 e817. 10.1016/j.chembiol.2022.02.004.

70. Austin, S.K., Dowd, K.A., Shrestha, B., Nelson, C.A., Edeling, M.A., Johnson, S., Pierson, T.C., Diamond, M.S., and Fremont, D.H. (2012). Structural basis of differential neutralization of DENV-1 genotypes by an antibody that recognizes a cryptic epitope. PLoS Pathog 8, e1002930. 10.1371/journal.ppat.1002930.

71. Hayes, E.B., and Gubler, D.J. (2006). West Nile virus: epidemiology and clinical features of an emerging epidemic in the United States. Annu Rev Med 57, 181–194. 10.1146/annurev.med.57.121304.131418.

72. Sejvar, J.J. (2014). Clinical manifestations and outcomes of West Nile virus infection. Viruses 6, 606–623. 10.3390/v6020606.

73. Angelou, A., Pappa, A., Markov, P.V., Gewehr, S., Stilianakis, N.I., and Kioutsioukis, I. (2025). Early warning system of the seasonal west nile virus infection risk in humans in northern greece, 2020-2024. Sci Rep 15, 7129. 10.1038/s41598-025-91996-9.

74. Meletis, E., Poulakida, I., Perlepe, G., Katsea, A., Pateras, K., Boutlas, S., Papadamou, G., Gourgoulianis, K., and Kostoulas, P. (2024). Early warning of potential epidemics: A pilot application of an early warning tool to data from the pulmonary clinic of the university hospital of Thessaly, Greece. J Infect Public Health 17, 401–405. 10.1016/j.jiph.2024.01.008.

75. Yun, S.I., and Lee, Y.M. (2014). Japanese encephalitis: the virus and vaccines. Hum Vaccin Immunother 10, 263–279. 10.4161/hv.26902.

76. Srivastava, K.S., Jeswani, V., Pal, N., Bohra, B., Vishwakarma, V., Bapat, A.A., Patnaik, Y.P., Khanna, N., and Shukla, R. (2023). Japanese Encephalitis Virus: An Update on the Potential Antivirals and Vaccines. Vaccines (Basel) 11. 10.3390/vaccines11040742.

77. Halstead, S.B., and Thomas, S.J. (2010). Japanese encephalitis: new options for active immunization. Clin Infect Dis 50, 1155–1164. 10.1086/651271.

78. Groen, K., Kuratli, R., Enkelmann, J., Fernbach, S., Wendel-Garcia, P.D., Staiger, W.I., Lejeune, M., Sauras-Colon, E., Roche-Campo, F., Filippidis, P., et al. (2025). Type I interferon autoantibody footprints reveal neutralizing mechanisms and allow inhibitory decoy design. J Exp Med 222. 10.1084/jem.20242039.

79. Cervantes Rincon, T., Kapoor, T., Keeffe, J.R., Simonelli, L., Hoffmann, H.H., Agudelo, M., Jurado, A., Peace, A., Lee, Y.E., Gazumyan, A., et al. (2024). Human antibodies in Mexico and Brazil neutralizing tick-borne flaviviruses. Cell Rep 43, 114298. 10.1016/j.celrep.2024.114298.

80. Meffre, E., Schaefer, A., Wardemann, H., Wilson, P., Davis, E., and Nussenzweig, M.C. (2004). Surrogate light chain expressing human peripheral B cells produce self-reactive antibodies. J Exp Med 199, 145–150. 10.1084/jem.20031550.

81. Klein, J.S., and Bjorkman, P.J. (2010). Few and far between: how HIV may be evading antibody avidity. PLoS Pathog 6, e1000908. 10.1371/journal.ppat.1000908.

82. Blight, K.J., McKeating, J.A., and Rice, C.M. (2002). Highly permissive cell lines for subgenomic and genomic hepatitis C virus RNA replication. J Virol 76, 13001–13014. 10.1128/jvi.76.24.13001-13014.2002.

83. Melnick, J.L., Paul, J.R., Riordan, J.T., Barnett, V.H., Goldblum, N., and Zabin, E. (1951). Isolation from human sera in Egypt of a virus apparently identical to West Nile virus. Proc Soc Exp Biol Med 77, 661–665. 10.3181/00379727-77-18884.

84. Rudolf, I., Bakonyi, T., Sebesta, O., Mendel, J., Pesko, J., Betasova, L., Blazejova, H., Venclikova, K., Strakova, P., Nowotny, N., and Hubalek, Z. (2014). West Nile virus lineage 2 isolated from Culex modestus mosquitoes in the Czech Republic, 2013: expansion of the European WNV endemic area to the North? Euro Surveill 19, 2–5. 10.2807/1560-7917.es2014.19.31.20867.

85. Serbia, O.G.o.t.R.o. (2019). Regulation on blood or blood component donors: 6/2019-132. https://www.pravno-informacioni-sistem.rs/SlGlasnikPortal/eli/rep/sgrs/ministarstva/pravilnik/2019/6/11.

86. Bianchini, F., Crivelli, V., Abernathy, M.E., Guerra, C., Palus, M., Muri, J., Marcotte, H., Piralla, A., Pedotti, M., De Gasparo, R., et al. (2023). Human neutralizing antibodies to cold linear epitopes and subdomain 1 of the SARS-CoV-2 spike glycoprotein. Sci Immunol 8, eade0958. 10.1126/sciimmunol.ade0958.

87. Robbiani, D.F., Gaebler, C., Muecksch, F., Lorenzi, J.C.C., Wang, Z., Cho, A., Agudelo, M., Barnes, C.O., Gazumyan, A., Finkin, S., et al. (2020). Convergent antibody responses to SARS-CoV-2 in convalescent individuals. Nature 584, 437–442. 10.1038/s41586-020-2456-9.

88. Foss, S., Sakya, S.A., Aguinagalde, L., Lustig, M., Shaughnessy, J., Cruz, A.R., Scheepmaker, L., Mathiesen, L., Ruso-Julve, F., Anthi, A.K., et al. (2024). Human IgG Fc-engineering for enhanced plasma half-life, mucosal distribution and killing of cancer cells and bacteria. Nat Commun 15, 2007. 10.1038/s41467-024-46321-9.

89. Lee, P.H., Huang, X.X., Teh, B.T., and Ng, L.M. (2019). TSA-CRAFT: A Free Software for Automatic and Robust Thermal Shift Assay Data Analysis. SLAS Discov 24, 606–612. 10.1177/2472555218823547.

90. Tiller, T., Meffre, E., Yurasov, S., Tsuiji, M., Nussenzweig, M.C., and Wardemann, H. (2008). Efficient generation of monoclonal antibodies from single human B cells by single cell RT-PCR and expression vector cloning. J Immunol Methods 329, 112–124. 10.1016/j.jim.2007.09.017.

91. von Boehmer, L., Liu, C., Ackerman, S., Gitlin, A.D., Wang, Q., Gazumyan, A., and Nussenzweig, M.C. (2016). Sequencing and cloning of antigen-specific antibodies from mouse memory B cells. Nat Protoc 11, 1908–1923. 10.1038/nprot.2016.102.

92. De Gasparo, R., Pedotti, M., Simonelli, L., Nickl, P., Muecksch, F., Cassaniti, I., Percivalle, E., Lorenzi, J.C.C., Mazzola, F., Magri, D., et al. (2021). Bispecific IgG neutralizes SARS-CoV-2 variants and prevents escape in mice. Nature 593, 424–428. 10.1038/s41586-021-03461-y.

93. Pierson, T.C., Sanchez, M.D., Puffer, B.A., Ahmed, A.A., Geiss, B.J., Valentine, L.E., Altamura, L.A., Diamond, M.S., and Doms, R.W. (2006). A rapid and quantitative assay for measuring antibody-mediated neutralization of West Nile virus infection. Virology 346, 53–65. 10.1016/j.virol.2005.10.030.

94. Schmidt, F., Weisblum, Y., Muecksch, F., Hoffmann, H.H., Michailidis, E., Lorenzi, J.C.C., Mendoza, P., Rutkowska, M., Bednarski, E., Gaebler, C., et al. (2020). Measuring SARS-CoV-2 neutralizing antibody activity using pseudotyped and chimeric viruses. J Exp Med 217. 10.1084/jem.20201181.

95. Esswein, S.R., Gristick, H.B., Jurado, A., Peace, A., Keeffe, J.R., Lee, Y.E., Voll, A.V., Saeed, M., Nussenzweig, M.C., Rice, C.M., et al. (2020). Structural basis for Zika envelope domain III recognition by a germline version of a recurrent neutralizing antibody. Proc Natl Acad Sci U S A 117, 9865–9875. 10.1073/pnas.1919269117.

96. Kabsch, W. (2010). Integration, scaling, space-group assignment and post-refinement. Acta Crystallogr D Biol Crystallogr 66, 133–144. 10.1107/S0907444909047374.

97. Agirre, J., Atanasova, M., Bagdonas, H., Ballard, C.B., Basle, A., Beilsten-Edmands, J., Borges, R.J., Brown, D.G., Burgos-Marmol, J.J., Berrisford, J.M., et al. (2023). The CCP4 suite: integrative software for macromolecular crystallography. Acta Crystallogr D Struct Biol 79, 449–461. 10.1107/S2059798323003595.

98. McCoy, A.J., Grosse-Kunstleve, R.W., Adams, P.D., Winn, M.D., Storoni, L.C., and Read, R.J. (2007). Phaser crystallographic software. J Appl Crystallogr 40, 658–674. 10.1107/S0021889807021206.

99. Zwart, P.H., Afonine, P.V., Grosse-Kunstleve, R.W., Hung, L.W., Ioerger, T.R., McCoy, A.J., McKee, E., Moriarty, N.W., Read, R.J., Sacchettini, J.C., et al. (2008). Automated structure solution with the PHENIX suite. Methods Mol Biol 426, 419–435. 10.1007/978-1-60327-058-8_28.

100. Emsley, P., and Cowtan, K. (2004). Coot: model-building tools for molecular graphics. Acta Crystallogr D Biol Crystallogr 60, 2126–2132. 10.1107/S0907444904019158.

101. Brochet, X., Lefranc, M.P., and Giudicelli, V. (2008). IMGT/V-QUEST: the highly customized and integrated system for IG and TR standardized V-J and V-D-J sequence analysis. Nucleic Acids Res 36, W503–508. 10.1093/nar/gkn316.

102. Meng, E.C., Goddard, T.D., Pettersen, E.F., Couch, G.S., Pearson, Z.J., Morris, J.H., and Ferrin, T.E. (2023). UCSF ChimeraX: Tools for structure building and analysis. Protein Sci 32, e4792. 10.1002/pro.4792.

103. Paxman, J.J., and Heras, B. (2017). Bioinformatics Tools and Resources for Analyzing Protein Structures. Methods Mol Biol 1549, 209–220. 10.1007/978-1-4939-6740-7_16.

104. Mouquet, H., Scharf, L., Euler, Z., Liu, Y., Eden, C., Scheid, J.F., Halper-Stromberg, A., Gnanapragasam, P.N., Spencer, D.I., Seaman, M.S., et al. (2012). Complex-type N-glycan recognition by potent broadly neutralizing HIV antibodies. Proc Natl Acad Sci U S A 109, E3268–3277. 10.1073/pnas.1217207109.

