## Supplementary figures for "Human antibodies against West Nile and related orthoflaviviruses"

### Figure S1

A

| Cohort | Age | Gender | | | |
|---|---|---|---|---|---|
| | | Male | | Female | |
| | | n | % | n | % |
| **WND** | 59.10 years (CI 95%: 55.145 – 63.055) | 45 | 65.21 | 24 | 34.78 |
| **WNF** | 42.33 years (CI 95%: 31.99 - 52.68) | 1 | 33.33 | 2 | 66.66 |

B

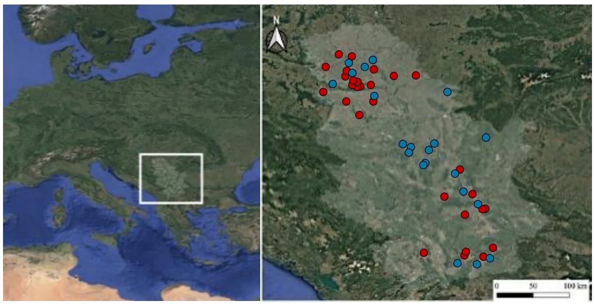

C

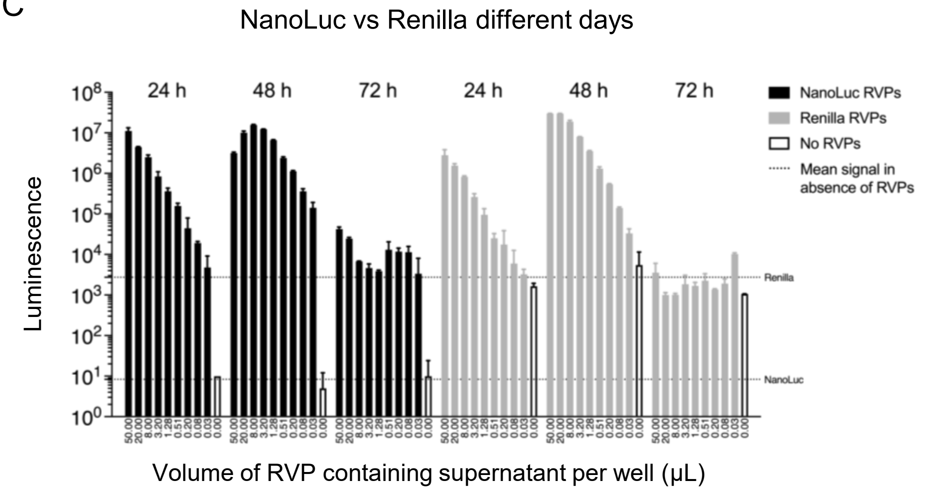


D

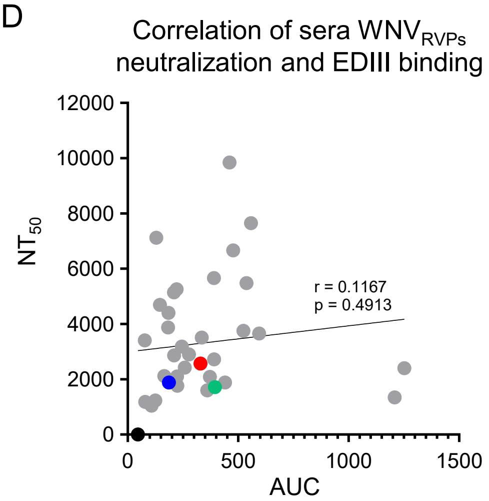


E F G H I J K L M N

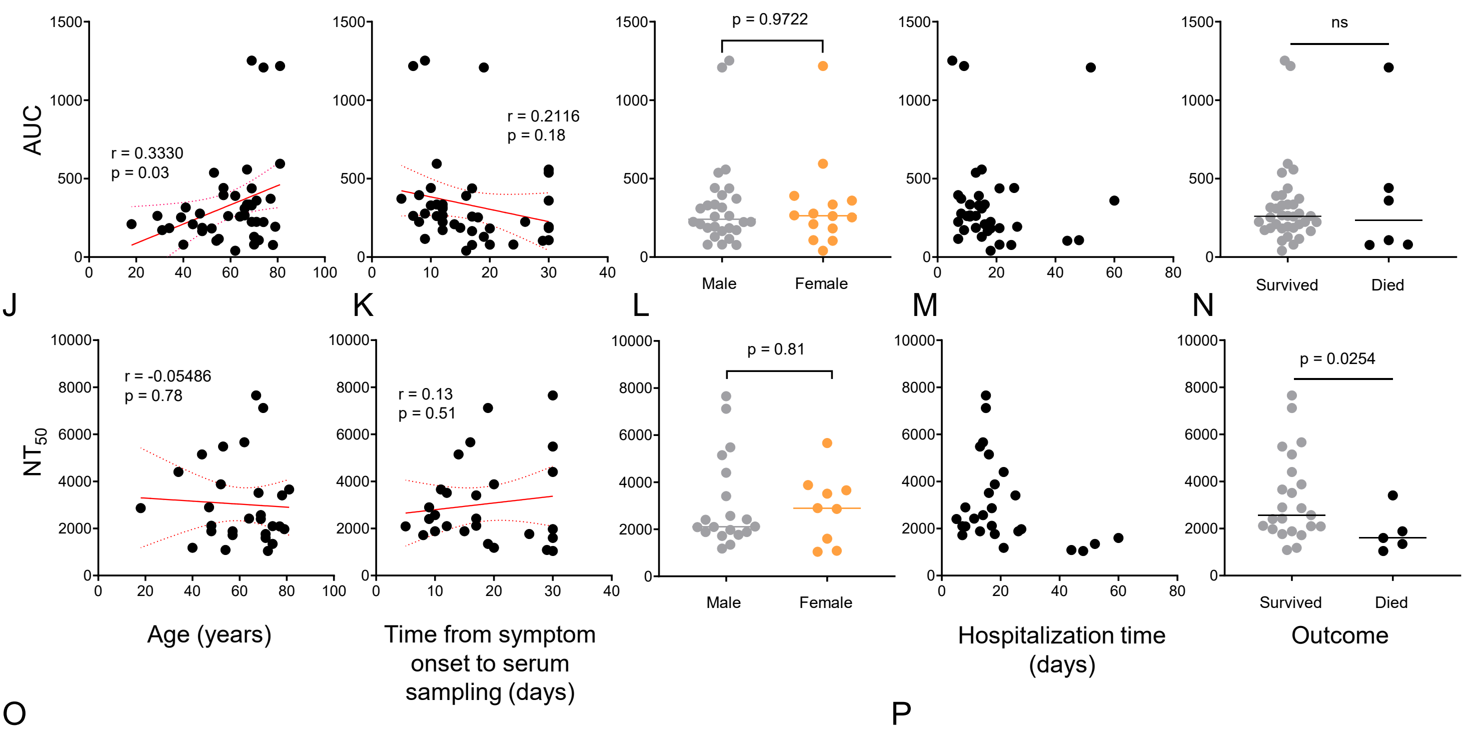


O

| anti-IFN-α2 autoantibodies | | | | | anti-IFN-ω autoantibodies | | | | |
|---|---|---|---|---|---|---|---|---|---|
| Age group | Serbian cohort | General population | p-value | Significantly higher? | Age group | Serbian cohort | General population | p-value | Significantly higher? |
| <50 | 0 / 8 (0%) | 8 / 2710 (0.30%) | — | No | <50 | 0 / 8 (0%) | 40 / 4446 (0.90%) | — | No |
| 50–60 | 2 / 7 (28.6%) | 3 / 1736 (0.17%) | < 0.001 | Yes | 50–60 | 1 / 7 (14.3%) | 13 / 1736 (0.75%) | 0.004 | Yes |
| 60–70 | 4 / 12 (33.3%) | 14 / 2475 (0.57%) | < 0.001 | Yes | 60–70 | 4 / 12 (33.3%) | 12 / 2475 (0.48%) | < 0.001 | Yes |
| 70–80 | 1 / 10 (10.0%) | 29 / 1790 (1.62%) | 0.15 | No | 70–80 | 1 / 10 (10.0%) | 29 / 1790 (1.62%) | 0.15 | No |
| 80–90 | 0 / 2 (0%) | 79 / 1580 (5.00%) | — | No | 80–90 | 0 / 2 (0%) | 56 / 1580 (3.54%) | — | No |
| **50–80** | **7 / 29 (24.1%)** | **46 / 6001 (0.766%)** | **< 0.001** | **Yes** | **50–80** | **6 / 29 (20.6%)** | **54 / 6001 (0.899%)** | **< 0.001** | **Yes** |
| P-values from two-sample, one-sided Fisher's exact tests | | | | | | | | | |
| "Significantly higher" defined as p < 0.05 | | | | | | | | | |
| "—" indicates no test due to zero events in the cohort | | | | | | | | | |

P

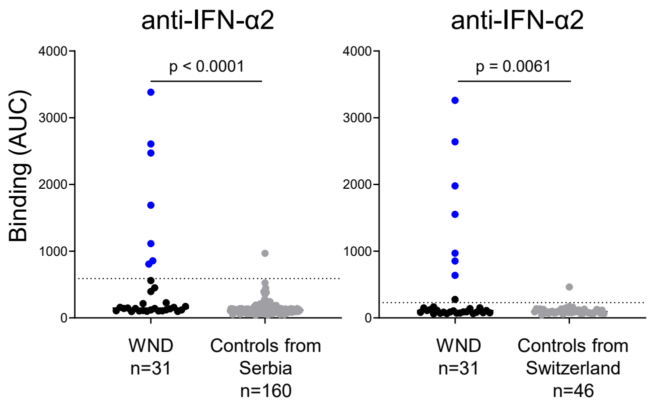

### Figure S2

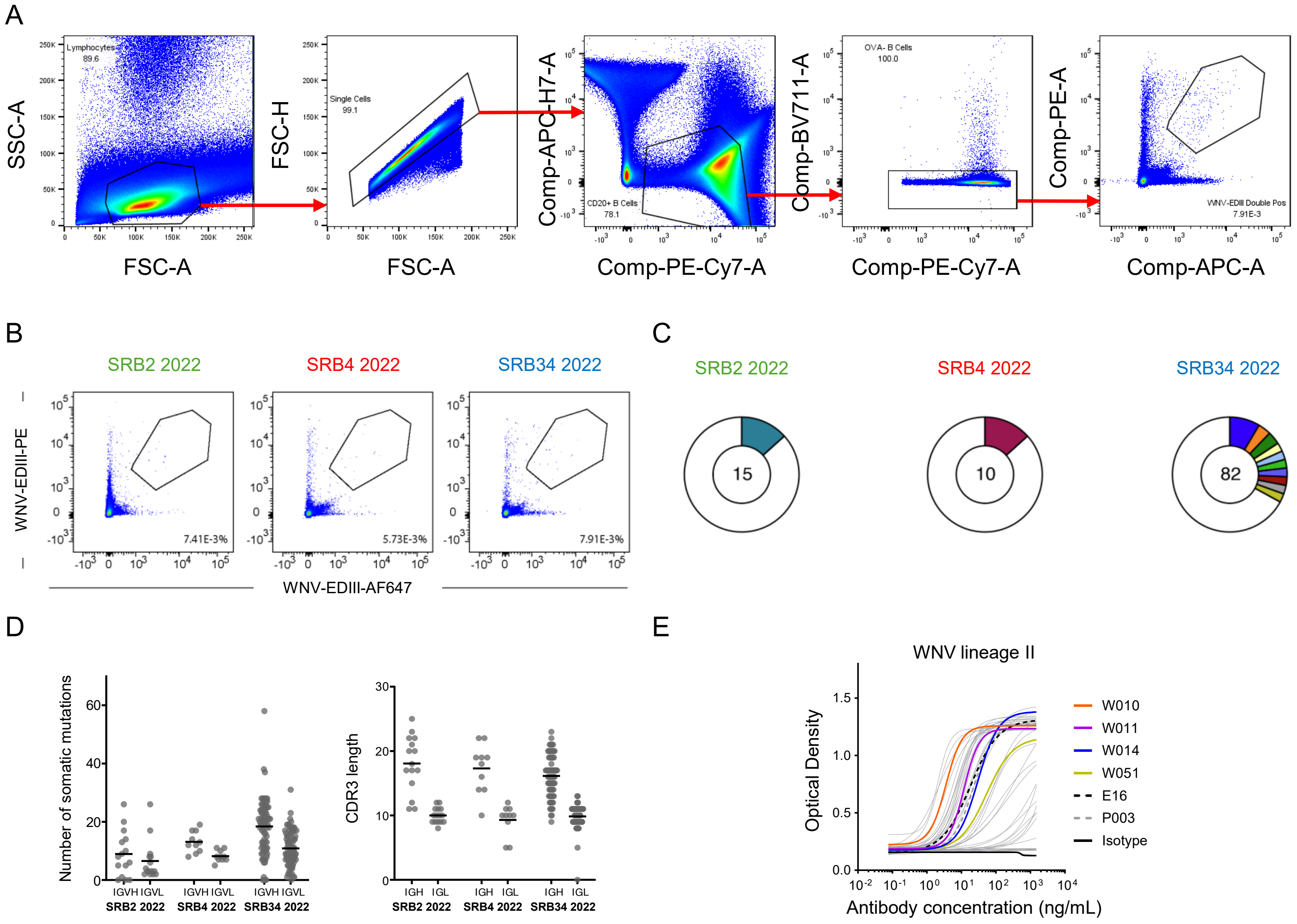

A
Lymphocytes 89.6
Single Cells 99.1
CD20+ B Cells 78.1
OVA- B Cells 100.0
WNV-EDIII Double Pos 7.91E-3
SSC-A
FSC-A
FSC-H
FSC-A
Comp-APC-H7-A
Comp-PE-Cy7-A
Comp-BV711-A
Comp-PE-Cy7-A
Comp-PE-A
Comp-APC-A
B
SRB2 2022
SRB4 2022
SRB34 2022
WNV-EDIII-PE
WNV-EDIII-AF647
7.41E-3%
5.73E-3%
7.91E-3%
C
SRB2 2022
SRB4 2022
SRB34 2022
15
10
82
D
Number of somatic mutations
IGVH IGVL
SRB2 2022
SRB4 2022
SRB34 2022
CDR3 length
IGH IGL
E
WNV lineage II
Optical Density
Antibody concentration (ng/mL)
W010
W011
W014
W051
E16
P003
Isotype

### Figure S3

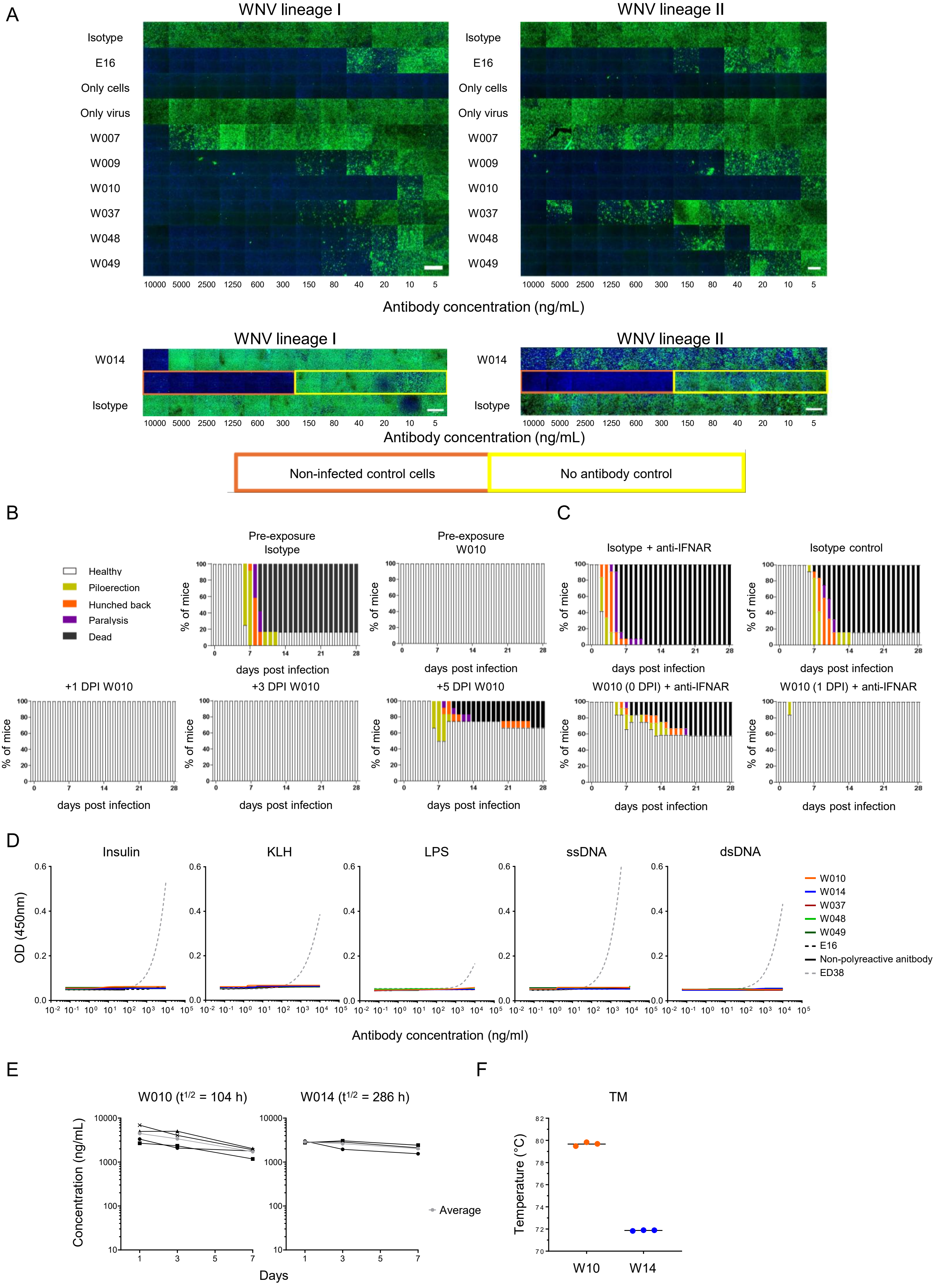

A
WNV lineage I
WNV lineage II
Isotype
E16
Only cells
Only virus
W007
W009
W010
W037
W048
W049
10000 5000 2500 1250 600 300 150 80 40 20 10 5
Antibody concentration (ng/mL)
W014
Isotype
Non-infected control cells
No antibody control
B
C
Healthy
Piloerection
Hunched back
Paralysis
Dead
Pre-exposure Isotype
Pre-exposure W010
Isotype + anti-IFNAR
Isotype control
+1 DPI W010
+3 DPI W010
+5 DPI W010
W010 (0 DPI) + anti-IFNAR
W010 (1 DPI) + anti-IFNAR
% of mice
days post infection
D
Insulin
KLH
LPS
ssDNA
dsDNA
OD (450nm)
W010
W014
W037
W048
W049
E16
Non-polyreactive anitbody
ED38
Antibody concentration (ng/ml)
E
W010 ($t^{1/2}$ = 104 h)
W014 ($t^{1/2}$ = 286 h)
Concentration (ng/mL)
Average
Days
F
TM
Temperature (°C)
W10
W14

### Figure S4

A

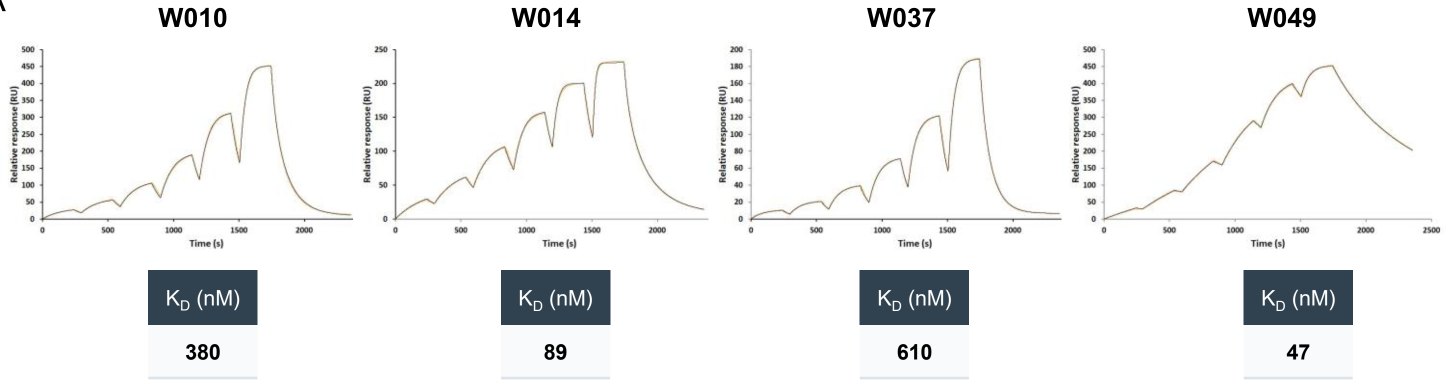


B

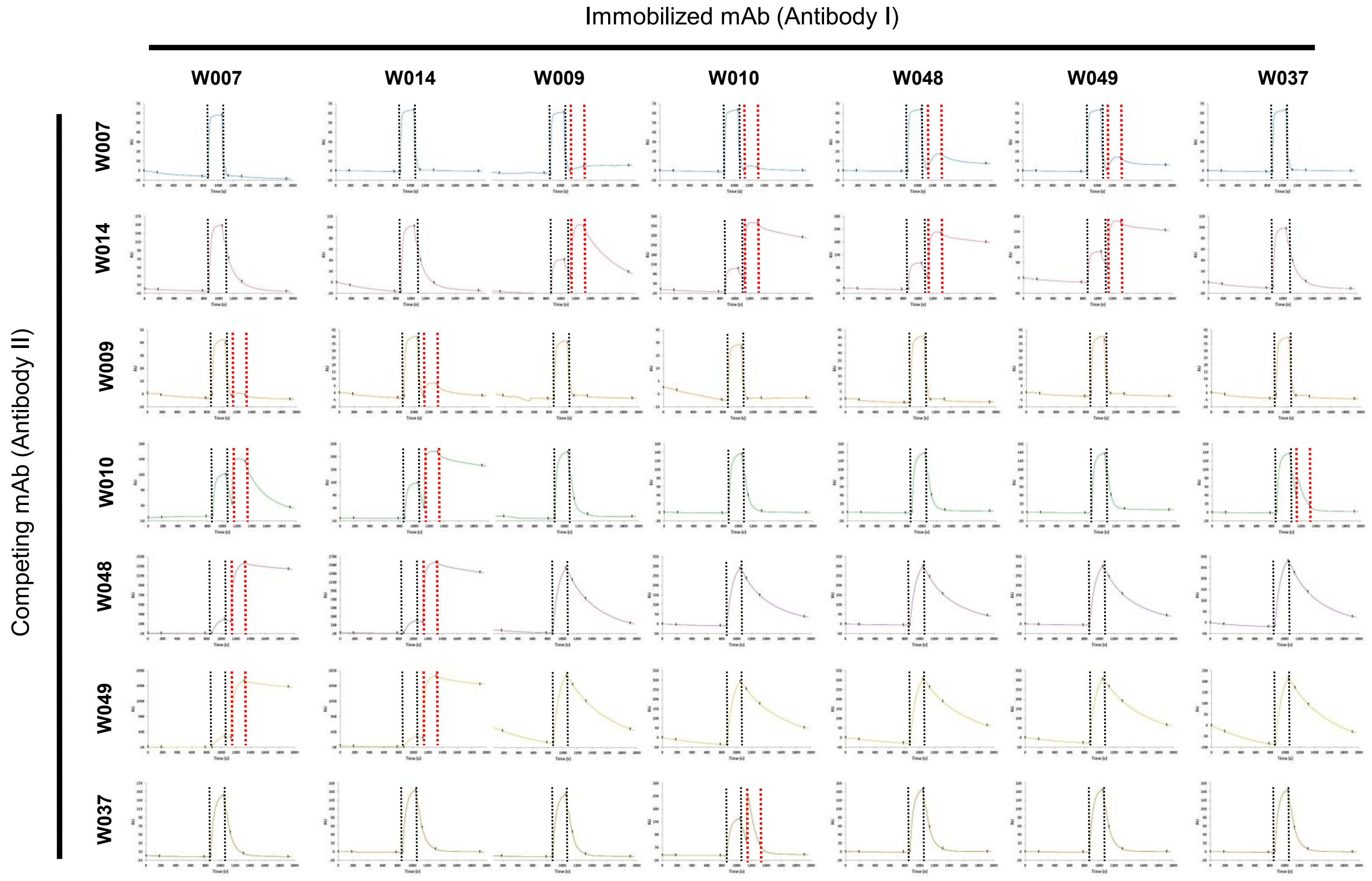


C

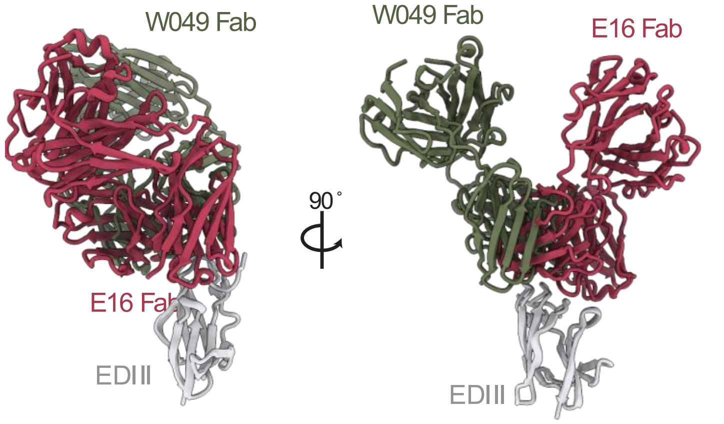


D

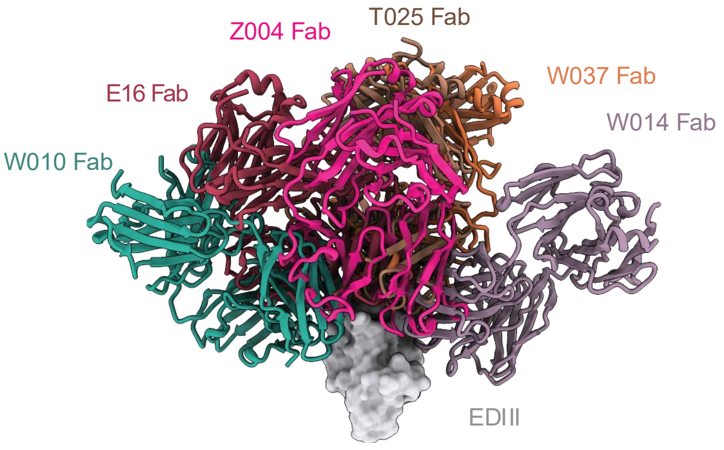

### Figure S5

A

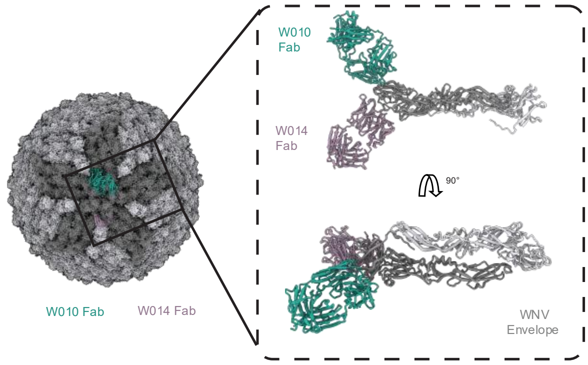


B

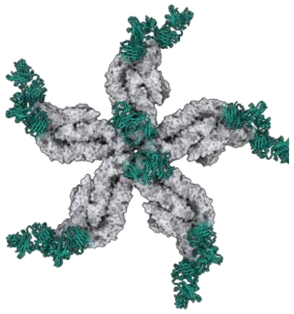

C

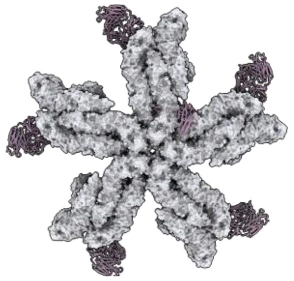

D

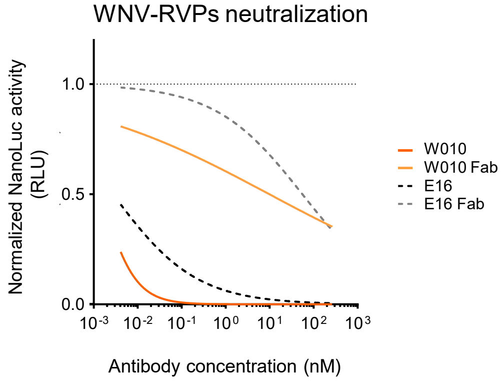


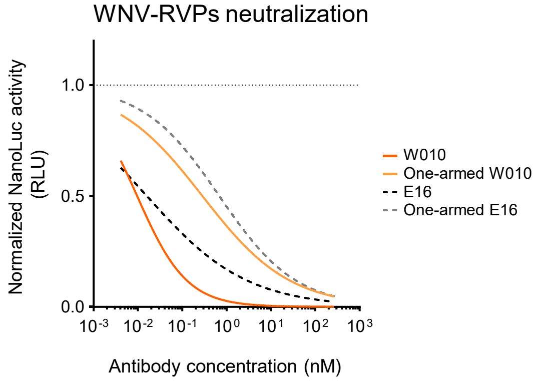


| Molecule | $IC_{50}$ value (nM) | MNR |
|---|---|---|
| W010 mAb | 0.00147 | 6673.5 |
| W010 Fab | 9.81 | |
| E16 mAb | 0.00273 | 20315.01 |
| E16 Fab | 55.46 | |

| Molecule | $IC_{50}$ value (nM) | MNR |
|---|---|---|
| W010 mAb | 0.009587 | 29.66 |
| One-armed W010 | 0.2844 | |
| E16 mAb | 0.01586 | 43.12 |
| One-armed E16 | 0.6840 | |

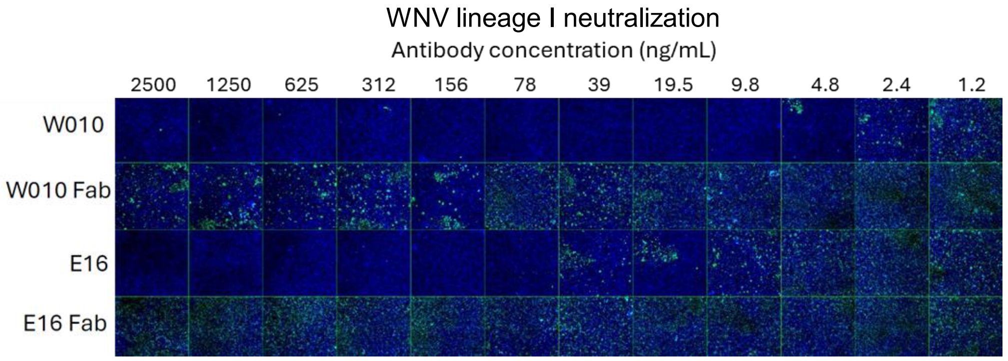


E

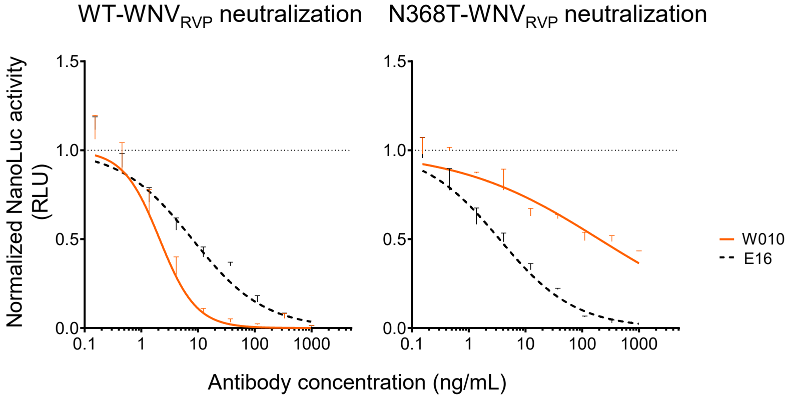

### Figure S6

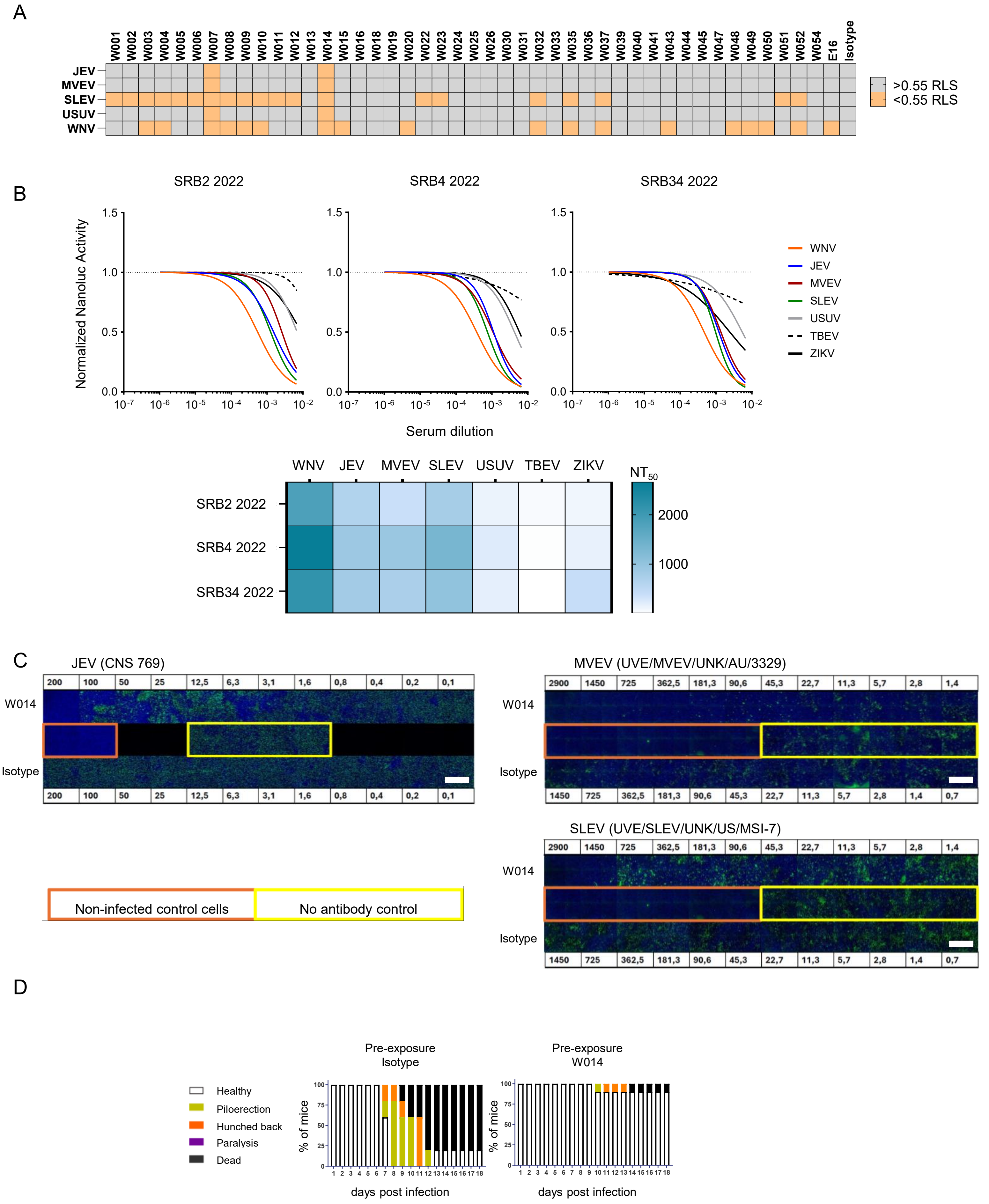
A
JEV
MVEV
SLEV
USUV
WNV
W001 W002 W003 W004 W005 W006 W007 W008 W009 W010 W011 W012 W013 W014 W015 W016 W018 W019 W020 W022 W023 W024 W025 W026 W030 W031 W032 W033 W035 W036 W037 W039 W040 W041 W043 W044 W045 W047 W048 W049 W050 W051 W052 W054 E16 Isotype
>0.55 RLS
<0.55 RLS
B
SRB2 2022
SRB4 2022
SRB34 2022
Normalized Nanoluc Activity
Serum dilution
WNV
JEV
MVEV
SLEV
USUV
TBEV
ZIKV
NT50
2000
1000
C
JEV (CNS 769)
MVEV (UVE/MVEV/UNK/AU/3329)
SLEV (UVE/SLEV/UNK/US/MSI-7)
W014
Isotype
Non-infected control cells
No antibody control
D
Healthy
Piloerection
Hunched back
Paralysis
Dead
Pre-exposure Isotype
Pre-exposure W014
% of mice
days post infection
