## Supplementary material for "Human antibodies against West Nile and related orthoflaviviruses": Table S1

**Table S1. Characteristics of study participants (n=72).**

| Participant ID | Gender (M/F) | Age at diagnosis (years) | Residency (place) | Suspected disease (WNF/WND) | Clinical diagnosis | High serum neutralization in the screening (yes/no) | Anti-type I IFN neutralization (IFNα2, IFNω, both) |
|---|---|---|---|---|---|---|---|
| SRB1 2022 | M | 57 | Kragujevac | WND | Viral meningoencephalitis (WNV) | yes | both |
| SRB2 2022 | M | 57 | Kragujevac | WND | Viral meningitis (WNV) | yes | n.d. |
| SRB3 2022 | M | 77 | Aranđelovac | WND | Viral meningoencephalitis (WNV) | yes | n.d. |
| SRB4 2022 | M | 69 | Topola | WND | Viral meningoencephalitis (WNV) | yes | n.d. |
| SRB5 2022 | F | 66 | Velika Plana | WND | Viral meningoencephalitis (WNV) | no | n.d. |
| SRB6 2022 | F | 68 | Aranđelovac | WND | Viral meningoencephalitis (WNV) | yes | n.d. |
| SRB7 2022 | M | 67 | Topola | WND | Viral meningoencephalitis (WNV) | no | n.d. |
| SRB8 2022 | M | 66 | Kragujevac | WND | Viral meningoencephalitis (WNV) | no | n.d. |
| SRB9 2022 | F | 59 | Rača | WND | Viral meningoencephalitis (WNV) | no | n.d. |
| SRB10 2022 | F | 81 | Kragujevac | WND | Viral meningoencephalitis (WNV) | no | n.d. |
| SRB11 2022 | M | 31 | Topola | WND | Viral meningoencephalitis (WNV) | no | n.d. |
| SRB12 2022 | M | 69 | Paraćin | WND | Viral meningoencephalitis (WNV) | no | n.d. |
| SRB14 2022 | M | 50 | Kragujevac | WND | Viral meningoencephalitis | yes | NT |
| SRB15 2022 | M | 64 | Kragujevac | WND | Viral meningoencephalitis (WNV) | yes | both |
| SRB16 2022 | F | 47 | Kragujevac | WNF | West Nile fever | yes | NT |
| SRB17 2022 | M | 29 | Aranđelovac | WND | Viral meningoencephalitis (WNV) | no | n.d. |
| SRB18 2022 | M | 55 | Topola | WND | Viral meningoencephalitis (WNV) | no | n.d. |
| SRB21 2022 | M | 64 | Donji Ljubeš | WND | Status febrilis, encephalopathia | yes | NT |
| SRB22 2022 | M | 49 | Vrtište | WND | Viral encephalitis | no | NT |
| SRB23 2022 | F | 39 | Niš | WNF | West Nile fever | no | NT |
| SRB24 2022 | M | 41 | Niš | WNF | West Nile fever | no | NT |
| SRB25 2022 | M | 78 | Gnjilane | WND | Viral encephalitis | no | NT |
| SRB26 2022 | F | 63 | Majdanpek | WND | Viral encephalitis | no | NT |
| SRB27 2022 | M | 73 | Vranjska banja | WND | Viral encephalitis | no | NT |
| SRB28 2022 | M | 66 | Bujanovac | WND | Viral encephalitis | no | NT |
| SRB29 2022 | M | 68 | Bačka Palanka | WND | Viral encephalitis | no | NT |
| SRB30 2022 | F | 76 | Bečej | WND | Viral encephalitis | no | NT |
| SRB31 2022 | F | 54 | Nadalj | WND | Viral encephalitis (WNV) | yes | n.d. |
| SRB32 2022 | M | 63 | Stepanovićevo | WND | Viral encephalitis | no | NT |
| SRB33 2022 | M | 48 | Novi Sad | WND | Viral encephalitis (WNV) | yes | n.d. |
| SRB34 2022 | M | 48 | Inđija | WND | Viral encephalitis (WNV) | yes | n.d. |
| SRB35 2022 | F | 62 | Novi Sad | WND | ncephalitis (WNV) and non-Hodgkin lyn | no | n.d. |
| SRB36 2022 | M | 40 | Vrbas | WND | Viral encephalitis (WNV) | yes | n.d. |
| SRB37 2022 | M | 29 | Novi Sad | WND | Viral encephalitis | no | NT |
| SRB38 2022 | F | 72 | Golubinci | WND | Viral encephalitis (WNV) | yes | n.d. |
| SRB1 2023 | M | 53 | Grabovci | WND | WNV encephalitis | yes | IFNa2 |
| SRB2 2023 | F | 62 | Sečanj | WND | WNV encephalitis | yes | n.d. |
| SRB3 2023 | F | 52 | Šid | WND | WNV encephalitis | yes | n.d. |
| SRB4 2023 | M | 70 | Odžaci | WND | WNV encephalitis | no | n.d. |
| SRB5 2023 | M | 44 | Čurug | WND | WNV encephalitis | yes | n.d. |
| SRB6 2023 | F | 81 | Sremska Kamenica | WND | WNV encephalitis | yes | n.d. |
| SRB7 2023 | M | 67 | Ravno Selo | WND | WNV encephalitis | yes | both |
| SRB8 2023 | M | 88 | Futog | WND | Encephalitis viralis non-specificata | yes | NT |
| SRB9 2023 | M | 79 | Ledinci | WND | WNV encephalitis | yes | n.d. |
| SRB10 2023 | M | 74 | Golubinci | WND | WNV encephalitis | yes | n.d. |
| SRB11 2023 | M | 71 | Novi Sad | WND | WNV encephalitis | yes | n.d. |
| SRB12 2023 | M | 70 | Novi Sad | WND | WNV encephalitis | yes | n.d. |
| SRB13 2023 | M | 34 | Feketić | WND | Meningoencephalitis WNV | yes | n.d. |
| SRB14 2023 | M | 69 | Prokuplje | WND | WNV encephalitis | yes | both |
| SRB15 2023 | F | 18 | Nova Crvenka | WND | Meningoencephalitis WNV | yes | n.d. |
| SRB16 2023 | M | 78 | Kovilj | WND | Encephalitis acuta | yes | NT |
| SRB17 2023 | M | 64 | Zrenjanin | WND | Encephalitis viralis non-specificata | yes | NT |
| SRB18 2023 | M | 62 | Rumenka | WND | Encephalitis viralis non-specificata | yes | NT |
| SRB19 2023 | M | 72 | Bački Petrovac | WND | Encephalomyelitis | yes | NT |
| SRB20 2023 | M | 78 | Šid | WND | WNV encephalitis | yes | n.d. |
| SRB21 2023 | M | 74 | Futog | WND | WNV encephalitis | yes | both |
| SRB22 2023 | M | 69 | Sremska Mitrovica | WND | WNV encephalitis | no | both |
| SRB23 2023 | F | 71 | Sremska Kamenica | WND | WNV encephalitis | yes | n.d. |
| SRB24 2023 | F | 35 | Niš | WND | Encephalitis viralis non-specificata | no | NT |
| SRB25 2023 | F | 75 | Niš | WND | Encephalitis viralis non-specificata | no | NT |
| SRB26 2023 | F | 55 | Vranje | WND | Encephalitis viralis non-specificata | no | NT |
| SRB27 2023 | F | 49 | Niš | WND | Encephalitis viralis non-specificata | no | NT |
| SRB28 2023 | M | 24 | Kostadince | WND | Encephalitis viralis non-specificata | no | NT |
| SRB29 2023 | F | 64 | Sastavar | WND | Encephalitis viralis non-specificata | no | NT |
| SRB30 2023 | M | 77 | Niš | WND | Meningoencephalitis acuta | no | NT |
| SRB31 2023 | M | 30 | Glogovac | WND | Encephalitis viralis non-specificata | no | NT |
| SRB32 2023 | F | 59 | Niš | WND | Encephalopathia | no | NT |
| SRB33 2023 | F | 28 | Niš | WND | Encephalitis viralis non-specificata | no | NT |
| SRB34 2023 | F | 28 | Aleksinac | WND | Encephalitis viralis non-specificata | yes | NT |
| SRB35 2023 | M | 18 | Vladičin Han | WND | Encephalitis viralis non-specificata | no | NT |
| SRB36 2023 | F | 61 | Doljane | WND | Encephalitis viralis non-specificata | no | NT |
| SRB37 2023 | F | 57 | Popovac | WND | Encephalitis viralis non-specificata | no | NT |

NT = Not Tested
n.d. = non detectable
ND = no data

| Time from symptom onset to serum sampling (days) | Duration of hospitalization (days) | Outcome (survived/died) | Clinical Center |
|---|---|---|---|
| 10 | 26 | Died | Clinical Center Kragujevac |
| 8 | 7 | Survived | Clinical Center Kragujevac |
| 5 | 8 | Survived | Clinical Center Kragujevac |
| 10 | 14 | Survived | Clinical Center Kragujevac |
| 12 | 11 | Survived | Clinical Center Kragujevac |
| 12 | 16 | Survived | Clinical Center Kragujevac |
| 11 | 11 | Survived | Clinical Center Kragujevac |
| 12 | 15 | Survived | Clinical Center Kragujevac |
| 11 | 10 | Survived | Clinical Center Kragujevac |
| 7 | 9 | Survived | Clinical Center Kragujevac |
| 12 | 9 | Survived | Clinical Center Kragujevac |
| 8 | 7 | Survived | Clinical Center Kragujevac |
| 9 | 15 | Survived | Clinical Center Kragujevac |
| 17 | 11 | Survived | Clinical Center Kragujevac |
| 9 | 8 | Survived | Clinical Center Kragujevac |
| 7 | 13 | Survived | Clinical Center Kragujevac |
| 9 | 7 | Survived | Clinical Center Kragujevac |
| 12 | 12 | Survived | Clinical Center Niš |
| 11 | ND | ND | Clinical Center Niš |
| 18 | ND | Survived | Clinical Center Niš |
| 19 | ND | Survived | Clinical Center Niš |
| 5 | ND | ND | Clinical Center Niš |
| 4 | ND | ND | Clinical Center Niš |
| 6 | ND | ND | Clinical Center Niš |
| 5 | ND | ND | Clinical Center Niš |
| 4 | 10 | Survived | Clinical Center Novi Sad |
| 9 | ND | ND | Clinical Center Novi Sad |
| 29 | 44 | Survived | Clinical Center Novi Sad |
| 7 | ND | ND | Clinical Center Novi Sad |
| 17 | 17 | Survived | Clinical Center Novi Sad |
| 15 | 13 | Survived | Clinical Center Novi Sad |
| 16 | 18 | Survived | Clinical Center Novi Sad |
| 20 | 21 | Survived | Clinical Center Novi Sad |
| 17 | ND | ND | Clinical Center Novi Sad |
| 30 | 48 | Died | Clinical Center Novi Sad |
| 30 | 13 | Survived | Clinical Center Novi Sad |
| 16 | 14 | Survived | Clinical Center Novi Sad |
| 20 | 18 | Survived | Clinical Center Novi Sad |
| 24 | 21 | Died | Clinical Center Novi Sad |
| 14 | 16 | Survived | Clinical Center Novi Sad |
| 11 | ND | Survived | Clinical Center Novi Sad |
| 30 | 15 | Survived | Clinical Center Novi Sad |
| ND | 14 | Survived | Clinical Center Novi Sad |
| 30 | 27 | Survived | Clinical Center Novi Sad |
| 12 | 7 | Survived | Clinical Center Novi Sad |
| 26 | 18 | Survived | Clinical Center Novi Sad |
| 19 | 15 | Survived | Clinical Center Novi Sad |
| 30 | 21 | Survived | Clinical Center Novi Sad |
| 9 | 5 | Survived | Clinical Center Novi Sad |
| ND | 17 | Survived | Clinical Center Novi Sad |
| 15 | 17 | Survived | Clinical Center Novi Sad |
| 20 | 20 | Survived | Clinical Center Novi Sad |
| 25 | 28 | Survived | Clinical Center Novi Sad |
| 16 | 16 | Survived | Clinical Center Novi Sad |
| 17 | 25 | Died | Clinical Center Novi Sad |
| 19 | 52 | Died | Clinical Center Novi Sad |
| 17 | 21 | Survived | Clinical Center Novi Sad |
| 30 | 60 | Died | Clinical Center Novi Sad |
| 5 | ND | ND | Clinical Center Niš |
| 5 | ND | ND | Clinical Center Niš |
| 5 | ND | ND | Clinical Center Niš |
| 8 | ND | ND | Clinical Center Niš |
| 4 | ND | ND | Clinical Center Niš |
| 3 | ND | ND | Clinical Center Niš |
| 5 | ND | ND | Clinical Center Niš |
| 5 | ND | ND | Clinical Center Niš |
| 2 | ND | ND | Clinical Center Niš |
| 5 | ND | ND | Clinical Center Niš |
| 8 | 20 | Survived | Clinical Center Niš |
| 10 | ND | ND | Clinical Center Niš |
| 7 | ND | ND | Clinical Center Niš |
| 10 | ND | ND | Clinical Center Niš |
