## Supplementary material for "Human antibodies against West Nile and related orthoflaviviruses": Table S2

**Table S3. Effective and inhibitory concentrations of recombinantly expressed monoclonal antibodies**

| | $EC_{50}$ (ng/mL) | | | | | |
|---|---|---|---|---|---|---|
| | WNV lin I - EDIII | WNV lin II - EDIII | JEV - EDIII | MVEV - EDIII | SLEV - EDIII | USUV - EDIII |
| W001 | | | | | | |
| W002 | | | | | | |
| W003 | | | | | | |
| W004 | 132 | 83.6 | | | | |
| W005 | 2.325 | 2.092 | | | | |
| W006 | | | | | | |
| W007 | | | | | | |
| W008 | 20.6 | 10.66 | | | | |
| W009 | 4.281 | 2.204 | | | | |
| W010 | 4.312 | 3.532 | | | | |
| W011 | 9.398 | 11.88 | 17.27 | 203.1 | | 17.89 |
| W012 | 6.792 | 3.747 | | | | |
| W013 | | | | | | |
| W014 | 42.22 | 30.06 | 31.41 | | 15.94 | 17.84 |
| W015 | 14.42 | 7.982 | | | | |
| W016 | 9.385 | 8.695 | | | | |
| W018 | 13.04 | 2.149 | | | | |
| W019 | 64.5 | 27.95 | | | | |
| W020 | | | | | | |
| W022 | 9.456 | 6.684 | | | | |
| W023 | 13.79 | 10.75 | | | | |
| W024 | | | | | | |
| W025 | 89.45 | 22.09 | | | | |
| W026 | | | | | | |
| W030 | | | | | | |
| W031 | | | | | | |
| W032 | | | | | | |
| W033 | | | | | | |
| W035 | 33.42 | 16.65 | | | | |
| W036 | | | | | | |
| W037 | 17.54 | 19.38 | | | | |
| W039 | | | | | | |
| W040 | 173.6 | 129 | | | | |
| W041 | | | | | | |
| W043 | 8.256 | 7.048 | | | | |
| W044 | 23.33 | 11.12 | | | | |
| W045 | 18.46 | 8.676 | | | | |
| W047 | 4.958 | 7.519 | | | | |
| W048 | 4.603 | 6.017 | | | | |
| W049 | 14.54 | 17.2 | | | | |
| W050 | 97.35 | 86.75 | | | | |
| W051 | 53.47 | 53.68 | | | 17.19 | |
| W052 | | | | | | |
| W054 | 13.18 | 13.82 | | | | |

Not-determined
Not-tested
Non-neutralizing

| $IC_{50}$ (ng/mL) - RVPs | | | | | $IC_{50}$ (ng/mL) - virus | |
|---|---|---|---|---|---|---|
| $WNV_{RVPs}$ lin I | $JEV_{RVPs}$ | $MVEV_{RVPs}$ | $SLEV_{RVPs}$ | $USUV_{RVPs}$ | WNV lin I | WNV lin II |
| 90.21 | 166.8 | 88.91 | 13.44 | 59.82 | 107.1 | 85.15 |
| 15 | | | | | | |
| 782.1 | | | | | 24.71 | 18.23 |
| 3.696 | | | | | 4.819 | 3.839 |
| 102.2 | 43.61 | 27.77 | 4.327 | 36.27 | 772.9 | 142.9 |
| 13.48 | | | | | | |
| 238.3 | | | | | | |
| 152.8 | | | | | | |
| 113.2 | | | | | | |
| | | | 140.2 | | | |
| 6.696 | | | | | 56.78 | 35.9 |
| 34.25 | | | | | 14.14 | 13.09 |
| 2.028 | | | | | 10.39 | 11.65 |
| 14.11 | | | | | | |
| 1347 | | 485.7 | 12.8 | | | |
| 212.1 | | | | | | |
