## Supplementary material for "Human antibodies against West Nile and related orthoflaviviruses": Table S3

Table S3. Crystallographic data processing and refinement statistics

| | | W010-WNV EDIII | W014-WNV EDIII | W037-WNV EDIII | W049-WNV EDIII |
|---|---|---|---|---|---|
| **PDB ID** | | 9ZRM | 9ZRN | 9ZRO | 9ZRP |
| **Data collection[a]** | | | | | |
| Space group | | P1 | P 1 21 1 | P 21 21 2 | P 21 21 21 |
| Unit cell (Å) | | 39.9 56.8 67.8 | 41.1 122.1 54.1 | 74.9 158.9 52.4 | 97.179 112.691 129.888 |
| α, β, γ (°) | | 66.5 89.4 70.2 | 90 106.331 90 | 90 90 90 | 90 90 90 |
| Wavelength (Å) | | 0.97946 | 0.97946 | 0.97946 | 0.97946 |
| Resolution (Å) | | 37.14 - 1.8 (1.864 - 1.8) | 39.54 - 2.1 (2.175 - 2.1) | 37.76 - 1.4 (1.45 - 1.4) | 39.55 - 2.5 (2.56 - 2.5) |
| Unique Reflections | | 42869 (4320) | 29372 (2825) | 122938 (11699) | 93009 (6498) |
| Completeness (%) | | 90.80 (91.79) | 98.35 (93.76) | 99.38 (95.62) | 97.86 (96.99) |
| Redundancy | | 3.9 (3.8) | 3.6 (3.1) | 6.7 (6.3) | 6.9 (7) |
| $CC_{1/2}$ (%) | | 99.7 (94.3) | 99.1 (51.7) | 99.9 (62.1) | 99.6 (48.7) |
| $\langle I/\sigma I \rangle$ | | 11.9 (3.8) | 5.42 (1.38) | 12.29 (2.13) | 8.7 (1.2) |
| Mosaicity (˚) | | 0.12 | 0.17 | 0.09 | 0.15 |
| $R_{merge}$ (%) | | 6.6 (30.3) | 8.579 (60.53) | 2.812 (38.43) | 14.7 (95.1) |
| $R_{pim}$ (%) | | 3.8 (17.9) | 8.579 (60.53) | 2.812 (38.43) | 9.1(63.5) |
| Wilson *B*-factor | | 17.6 | 28.09 | 13.75 | 55.18 |
| **Refinement and Validation** | | | | | |
| Resolution (Å) | | 37.14 - 1.8 | 39.54 - 2.1 | 37.76 - 1.4 | 39.55 - 2.5 |
| Number of atoms | | | | | |
| | Protein | 4016 | 4096 | 4016 | 8365 |
| | Ligand | 0 | 0 | 0 | |
| | Waters | 480 | 367 | 788 | 176 |
| $R_{work}/R_{free}$ (%) | | 17.7/21.6 | 19.8/25.1 | 18.3/21.1 | 20.5/26.2 |
| R.m.s. deviations | | | | | |
| | Bond lengths (Å) | 0.004 | 0.024 | 0.013 | 0.008 |
| | Bond angles (˚) | 0.78 | 1 | 1.3 | 0.98 |
| MolProbity score | | 0.98 | 1.46 | 1.06 | 1.78 |
| Clashscore (all atom) | | 1.88 | 3.9 | 1.76 | 4.75 |
| Poor rotamers (%) | | 0.44 | 1.3 | 0.88 | 1.64 |
| Ramachandran plot | | | | | |
| | Favored (%) | 97.89 | 96.6 | 97.3 | 94.52 |
| | Allowed (%) | 2.1 | 3.7 | 2.67 | 5.38 |
| | Disallowed (%) | 0 | 0 | 0 | 0.09 |
| Average *B*-factor (Å) | | 22.36 | 32.13 | 19.9 | 58.41 |

[a]Numbers in parentheses correspond to the highest resolution shell
